# The Norepinephrine Metabolite DOPEGAL Confers Locus Coeruleus Tau Vulnerability and Propagation via Asparagine Endopeptidase

**DOI:** 10.1101/688200

**Authors:** Seong Su Kang, Xia Liu, Eun Hee Ahn, Jie Xiang, Fredric P. Manfredsson, Xifei Yang, Hongbo R. Luo, L. Cameron Liles, David Weinshenker, Keqiang Ye

## Abstract

Aberrant Tau inclusions in the locus coeruleus (LC) are the earliest detectable Alzheimer’s disease (AD)-like neuropathology in the human brain; however, why LC neurons are selectively vulnerable to developing early Tau pathology and degenerating later in disease and whether the LC might seed the stereotypical spread of Tau pathology to the rest of the brain remain unclear. Here we show that 3,4-dihydroxyphenylglycolaldehyde (DOPEGAL), which is produced exclusively in noradrenergic neurons by monoamine oxidase A (MAO-A) metabolism of norepinephrine (NE), activates asparagine endopeptidase (AEP) that cleaves Tau at residue N368 into aggregation- and propagation-prone forms, thereby leading to LC degeneration and the spread of Tau pathology. DOPEGAL triggers AEP-cleaved Tau aggregation *in vitro* and in intact cells, resulting in LC neurotoxicity and propagation of pathology to the forebrain. Thus, our findings reveal a novel molecular mechanism underlying the selective vulnerability of LC neurons in AD.

## Introduction

Alzheimer’s disease (AD) is an insidious, debilitating illness characterized by amyloid-β (Aβ) plaques and Tau neurofibrillary tangles (NFT), as well as neuronal cell death and neuroinflammation. Despite intense research into the pathologic hallmarks of AD, no disease-modifying therapies have emerged (1), and recent research efforts are focused on early AD detection and treatment. One area of interest in early AD involves the locus coeruleus (LC), the major brainstem noradrenergic nucleus that innervates and supplies norepinephrine (NE) to the forebrain and regulates attention, arousal, cognition, and stress responses (2). Hyperphosphorylated Tau, a “pretangle” form of the protein prone to aggregation, appears in the LC before any other area of the brain and is now considered the earliest detectable AD-like neuropathology (3–7). Moreover, degeneration of the LC is a ubiquitous feature of mid- to late-stage disease, contributing to the progression of forebrain pathology and cognitive impairment (6, 8–12), while restoration of noradrenergic signaling can ameliorate AD-like symptoms in animal models (13, 14). The progression of AD pathology follows a remarkably systematic pattern across individuals, and aberrant forms of Tau are capable of neuron-to-neuron propagation via a prion-like process of corruptive templating (15). Hyperphosphorylated Tau appears exclusively in the LC in Braak stage 0 (a-c), spreads to the entorhinal cortex in Braak stages I-II, then propagates to the hippocampus and frontal cortex in stages III-VI (3). Because hyperphosphorylated Tau can first be detected in the LC, and the LC sends dense projections to other vulnerable brain regions that display early Tau pathology (e.g., the transentorhinal cortex and hippocampus), some suggest that the LC might be one of the origins of Tau neuropathology in AD (4, 5). Although several theories have emerged to explain the selective vulnerability of LC neurons in AD, such as their unique anatomical, electrophysiological, morphological, and neurochemical characteristics (7, 16), the underlying molecular mechanisms have yet to be identified.

Asparagine endopeptidase (AEP; gene name *LGMN*) is a lysosomal endopeptidase that specifically cleaves its substrates after asparagine under conditions of acidosis (17). Recently, we reported that AEP acts as a delta-secretase that cleaves both amyloid precursor protein (APP) and Tau in an age-dependent manner in mouse and human AD brains (18, 19). AEP cuts APP at the N373 and N585 residues in the ectodomain and facilitates Aβ production by decreasing the steric hindrance for beta-secretase (BACE1). Depletion of AEP significantly reduces Aβ production and senile plaque formation in 5XFAD transgenic mice, leading to the restoration of synaptic activity and cognitive functions (19). In addition, AEP cleaves Tau at N255 and N368 and abolishes its microtubule assembly activity, resulting in hyperphosphorylation, aggregation, and NFT formation (18). Of note, the AEP-generated Tau 1-368 fragment is highly neurotoxic. Genetic deletion or pharmacological inhibition of AEP in transgenic mice that overexpress mutant human APP or aggregation-prone mutant human Tau prevents APP N585 and Tau N368 cleavage, ameliorates Aβ and Tau pathology, and reverses synaptic plasticity and cognitive deficits (18, 20). Thus, AEP plays a critical role in regulating Aβ and Tau pathogenesis. Most recently, we reported that AEP is activated by 3,4-dihydroxyphenylacetaldehyde (DOPAL), a highly toxic and oxidative dopamine (DA) metabolite, in the substantia nigra (21). Active AEP subsequently cleaves α-synuclein at N103 and promotes its aggregation and dopameringeric neuronal loss, leading to motor dysfunction in animal models of Parkinson’s disease (PD) (22). Analogous to toxic DA metabolites killing substantia nigra pars compacta neurons in PD, the noradrenergic phenotype of LC neurons itself may contribute to the vulnerability of these cells in AD. NE is converted into 3,4-dihydroxyphenylglycolaldehyde (DOPEGAL) by monoamine oxidase A (MAO-A) during the normal life cycle of catecholamine production and transmission, but is increased in the degenerating LC neurons of AD patients (23–25). Injection of DOPEGAL into rodent brains elicits adrenergic neuronal loss (26), and DOPEGAL toxicity is likely due to generation of free radicals and activation of mitochondrial permeability transition (27). The activation of AEP by DOPAL suggested to us that it might also be activated by DOPEGAL, thus triggering a cascade of events leading to Tau cleavage, hyperphosphorylation, aggregation, and neurotoxicity in LC neurons.

## Results

### DOPEGAL triggers Tau aggregation *in vitro* and in cells

To examine whether DOPEGAL directly modifies Tau, we incubated recombinant wild-type Tau with DOPEGAL for 24 h. We found DOPEGAL induced Tau aggregation, indicated by the accumulation of high molecular weight bands, in a concentration-dependent manner. As expected, AEP-truncated Tau N368 recombinant proteins were more prone to aggregation than full-length Tau (Figure 1A). To explore the specificity of DOPEGAL, we tested various catecholamines and their oxidative metabolites (500 μM). Compared to vehicle control, DOPEGAL strongly provoked Tau aggregation, while NE, DA, and DOPAL instead caused Tau protein degradation (Figure 1B). Time course and dose-response assays demonstrated that both DA and NE triggered dose- and time-dependent degradation of Tau, whereas DOPAL elicited Tau aggregation at lower doses (2.5-25 μM) but promoted degradation at higher doses (Supplemental Figure 1A-C). To investigate whether DOPEGAL elicits Tau fibrillization *in vitro*, we monitored the fluorescent intensity of recombinant Tau in the presence of Thioflavin T (ThioT) (30 μM). Compared to vehicle control, DOPEGAL strongly augmented Tau fibrillization over time, as revealed by the high-intensity ThioT fluorescence of Tau pre-formed fibrils (PFFs), while NE and DOPAL inhibited Tau fibrillization (Figure 1C). Transmission electronic microscopy (TEM) analysis confirmed that Tau PFFs formed tight fibrillar structures following DOPEGAL exposure (Supplemental Figure 1D).

**Figure 1.**
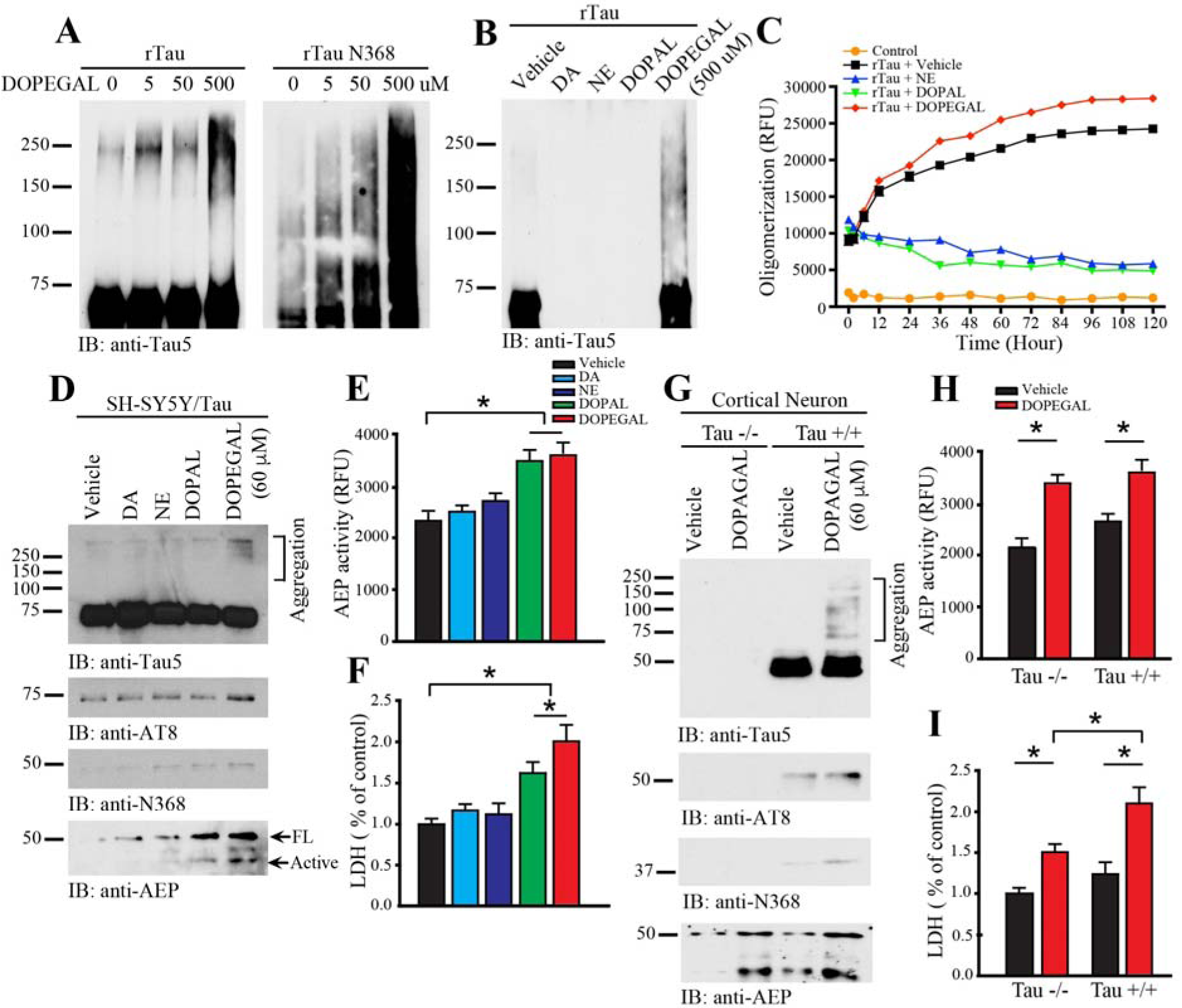
The NE metaboliote DOPEGAL triggers Tau aggregation *in vitro*. **A.** Recombinant Tau (rTau) or rTau N368 (1 μg) was incubated with DOPEGAL (0, 5, 50, or 500 μM) for 24 h, and aggregation was was detected by western blot using Tau5 antibody. **B.** rTau (1 μg) was incubated with vehicle, dopamine (DA), norepinephrine (NE), DOPAL, or DOPEGAL (500 μM) for 24 h, and aggregation was was detected by western blot using Tau5 antibody. **C.** rTau (1 μg) was incubated with vehicle or DOPEGAL, and ThioT assay was used to evaluate the kinetics of aggregation of Tau over 120 h. SH-SY5Y cells were infected with hTau virus, then exposed to DA, NE, DOPAL, or DOPEGAL (60 μM) for 24 h. **D.** Western blot analysis was conducted on cell lysates using antibodies against AEP and different forms of Tau. **E.** AEP activity and **F.** cell death were analyzed by enzymatic assay and LDH asssy, respectively. Data are shown as mean ± SEM. N=3 per group. * p<0.05. Primary cortical neurons were cultured from Tau +/+ and Tau −/− mice and exposed to vehicle or DOPEGAL (60 μM) for 24 h. **G.** Western blot analysis was conducted on cell lysates using antibodies against AEP and different forms of Tau. **H.** AEP activity and **I.** cell death were analyzed by enzymatic assay and LDH asssy, Data are shown as mean ± SEM. N=3 per group. * p<0.05.

To assess whether DOPEGAL provokes Tau aggregation in intact cells, we transfected noradrenergic-like SH-SY5Y cells with wild-type human Tau, followed by treatment with different catecholamines or their oxidative metabolites for 24 h. Immunoblotting revealed that only DOPEGAL treatment produced demonstrable Tau aggregation, which was in alignment with its hyperphosphorylation (AT8-positive) status (Figure 1D, top 2 panels). We next determined whether DOPEGAL, like DOPAL, activates AEP. We found DOPEGAL upregulated total AEP levels and its proteolytic activation, as well as the abundance of the AEP cleavage product Tau N368 (Figure 1D bottom 2 panels, Figure 1E), and induced SH-SY5Y cell death (Figure 1F). DOPAL was less effective in these measures, while NE and DA failed to activate AEP or cause toxicity. To ascertain whether Tau is required for DOPEGAL-elicited cell death, we prepared primary cortical neurons from neonatal wild-type (WT) and Tau knockout (Tau −/−) mice and treated them with DOPEGAL. As expected, DOPEGAL triggered demonstrable Tau hyperphosphorylation, aggregation, and N368 cleavage in WT, but not Tau −/−, neurons (Figure 1G). Importantly, while AEP was similarly activated by DOPEGAL in both WT and Tau −/− neurons (Figure 1H), its toxicity was significantly attenuated in Tau −/− neurons (Figure 1I), suggesting that Tau is necessary for the full expression of DOPEGAL-induced cell death.

### NE oxidation by MAO-A facilitates AEP-cleaved Tau N368 cytotoxicity

Oxidative deamination of NE by mitochondrial MAO-A generates DOPEGAL and H_2_O_2_, leading to oxidative stress (28, 29). To test whether MAO-mediated metabolism of NE results in oxidative stress and AEP activation, we transfected SH-SY5Y cells (30) with MAO-A or MAO-B. We found that MAO overexpression increased AEP enzymatic activity and Tau N368 cleavage (Supplemental Figure 2A-C), which was mimicked by H_2_O_2_ and prevented by the MAO-A inhibitor clorgyline, indicating the importance of oxidative stress (Supplemental Figure 2D-F). Interestingly, H_2_O_2_ also increased MAO-A expression, which was blunted by clorgyline. To evaluate the contribution of NE metabolism, we transfected SH-SY5Y cells with siRNA against dopamine β-hydroxylase (DBH), which is required for NE synthesis (31). DBH depletion reduced AEP activity, diminished Tau N368 cleavage, and partially ameliorated the deleterious effects of H_2_O_2_ (Supplemental Figure 2G-H). Combined, these results suggest that oxidative metabolism of NE to DOPEGAL by MAO activates AEP and Tau N368 cleavage in catecholaminergic SH-SY5Y cells.

To explore the neurotoxicity of AEP-cleaved Tau, we infected SH-SY5Y cells with AAV encoding full-length wild-type Tau, P301S Tau, the N368 Tau fragment, or uncleavable P301S/N255/368A Tau, and determined cell death using TUNEL staining and LDH assay (Supplemental Figure 2I-L). Wild-type Tau promoted cell death, which was escalated by the AEP-cleaved Tau N368 fragment. Notably, N368 Tau displayed neurotoxicity comparable to Tau P301S, while prevention of AEP cleavage blunted Tau P301S toxicity. These results indicate that AEP cleavage contributes to Tau neurotoxicity.

To ascertain whether DOPEGAL regulates Tau neurotoxicity, we overexpressed MAO-A or MAO-B in SH-SY5Y cells in combination with AAV-Tau. TUNEL and LDH analysis revealed that both MAO-A and MAO-B overexpression significantly activated AEP, increased Tau N368 cleavage, and magnified Tau neurotoxicity as measured by TUNEL, LDH, and loss of the catecholaminergic marker tyrosine hydroxylase (TH) (Supplemental Figure 3). By contrast, preventing DOPEGAL production by siRNA knockdown of DBH attenuated AEP activation and Tau-triggered cell death. These findings reveal that a proportion of Tau neurotoxicity is NE-dependent.

Our data support a model whereby MAO-A oxidizes NE into DOPEGAL and elicits oxidative stress, triggering AEP activation and Tau N368 cleavage and cytotoxicity. To further explore this idea, we overexpressed MAO-A and MAO-B into SH-SY5Y cells and interrogated their roles in AEP activation in the presence of specific inhibitors. We confirmed that overexpression of either MAO-A or MAO-B substantially provoked AEP activation, and blockade of these enzymes by small molecular inhibitors attenuated AEP enzymatic activity, Tau hyperphosphorylation, and N368 cleavage (Supplemental Figure 4A-E). To test whether elevation of NE in SH-SY5Y cells facilitates MAO-A cytotoxicity, we transfected cells with MAO-A, followed by treatment with ascorbic acid, which stimulates NE biosynthesis (32), the DA and NE precursor L-3,4-dihydroxyphenylalanine (L-DOPA), or the synthetic NE precursor L-3,4-dihydroxyphenylserine (L-DOPS). While these compounds alone barely activated AEP, both L-DOPS and ascorbic acid significantly increased AEP activation, Tau N368 cleavage, and AT8 immunoreactivity in the presence of elevated MAO-A (Supplemental Figure 4F-G). Interestingly, both L-DOPA and L-DOPS, but not ascorbic acid, exhibited cytotoxicity on their own, which was facilitated by MAO-A overexpression (Supplemental Figure 4H). The limited cytotoxicity of ascorbic acid may be attributed to its antioxidant activity.

DOPEGAL is metabolized *in vivo* via reduction by aldehyde reductase (AR), and further oxidation by aldehyde dehydrogenase (ALDH). To determine whether accumulation of DOPEGAL via blockade of its metabolism augments its cytotoxicity, we treated SH-SY5Y cells with the AR inhibitor imirestat, the ALDH inhibitor daidzein, or both drugs in the presence or absence of MAO-A overexpression. While none of the treatments were effective on their own, imirestat and the mixture of both inhibitors augmented AEP activation, Tau N368 cleavage, and AT8 abundance in combination with MAO-A (Supplemental Figure 4I-J). LDH assay indicated that the inhibitor cocktail modestly stimulated cell death without MAO-A overexpression, while either Imirestat or Imirestat + daidzein escalated MAO-A cytotoxicity (Supplemental Figure 4K). These results suggest that AR is the major enzyme responsible for DOPEGAL metabolism in SH-SY5Y cells, and that preventing DOPEGAL metabolism is deleterious.

### Tau hyperphosphorylation and N368 cleavage are increased in the LC of human AD subjects and transgenic AD mouse models

Hyperphosphorylated Tau in the LC is among the first detectable AD pathologies in postmortem human brains, appearing before Tau pathology in the EC and often decades prior to cognitive symptoms (5). To explore whether Tau N368 cleavage by AEP correlates with Tau hyperphosphorylation, we conducted immunofluorescence (IF) staining in the LC of postmortem human AD brains and different ages of transgenic AD mouse models. TH was used as a marker for noradrenergic LC neurons. AT8 and Tau N368 immunoreactivity were absent from WT mouse brains but appeared and accumulated together in a time-dependent manner in the LC of both 3xTg mice and Tau P301S mice. Compared to healthy controls, LC sections from AD subjects exhibited robust AT8 and Tau N368 signals (Figure 2A). These results indicate that AEP is temporally activated in the LC, cleaving Tau N368 and triggering its hyperphosphorylation. To investigate whether the hyperphosphorylated Tau was aggregated, we performed IF co-staining with AT8 or Tau N368 and Thioflavin S (ThioS). AT8/N368-positive Tau was also ThioS-positive in the LC region of 12-month-old 3xTg mice, 6-month-old Tau P301S mice, and human AD subjects (Figure 2B). IF co-staining also showed that AEP fluorescence intensity was elevated in the LC of both 3xTg and Tau P301S mice compared to WT controls, similar to human AD brains (Figure 2C). These findings demonstrate that activated AEP is abundant in the diseased LC, where it cleaves Tau N368, triggering its hyperphosphorylation and aggregation.

**Figure 2.**
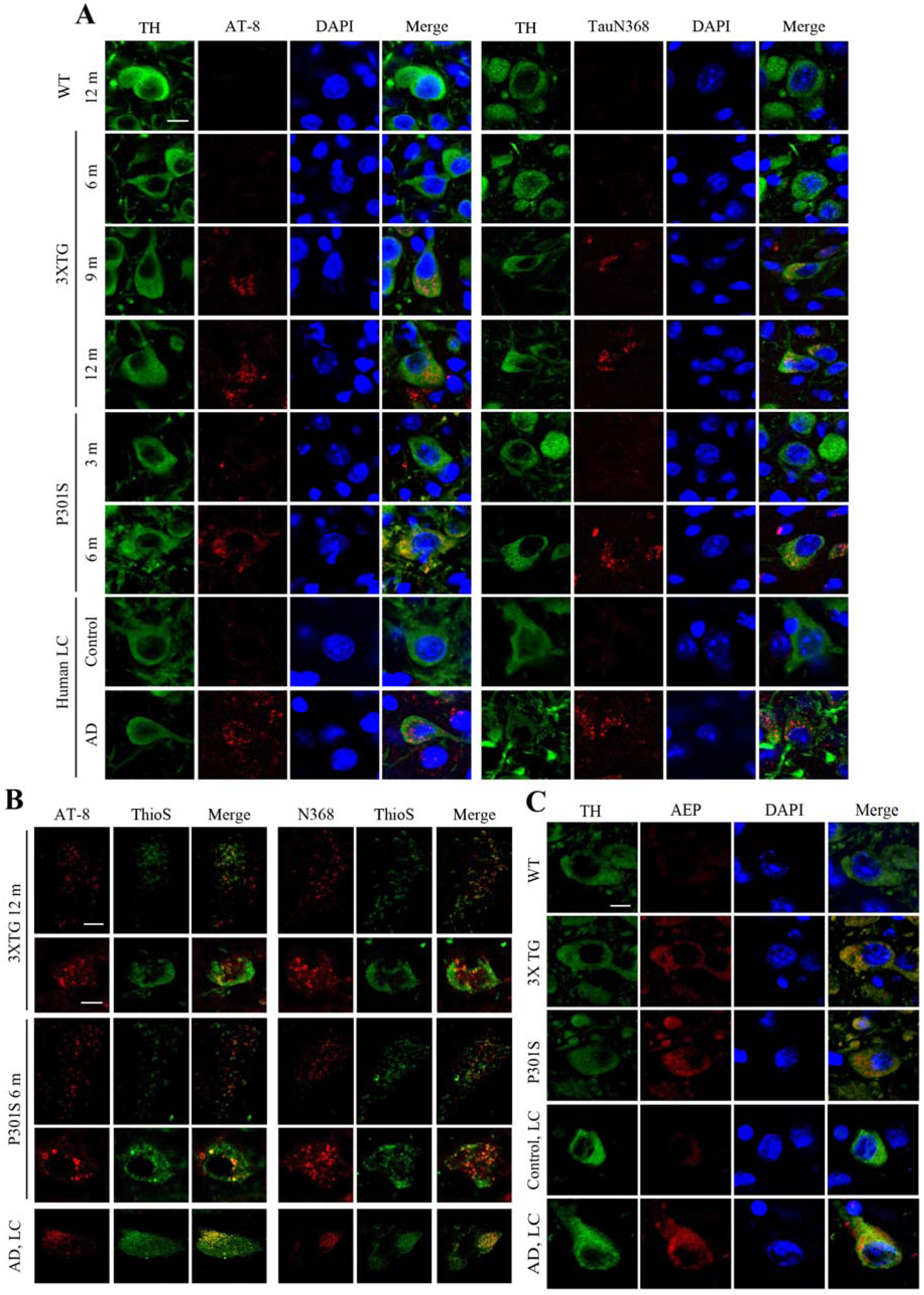
Tau is cleaved at N368 and hyperphosphorylated in the LC region of transgenic AD mouse models and human postmortem AD brains. Wild-type (WT), 3xTg, and P301S mouse brains from various ages, as well as postmortem human control and AD brains, were examined for Tau pathology in the LC by immunohistochemistry. **A.** Representative images of staining for tyrosine hydroxylase (TH; green), AT-8 or Tau N368 (red), and DAPI (blue). Scale bar = 20 μm. **B.** Representative images of staining for AT-8 or Tau N368 (red) and ThioS (green). Scale bar of top panel is 100 μm and second panel is 20 μm**. C.** Representative images of staining for TH (green), AEP (red), and DAPI (blue). Scale bar = 20 μm.

### NE is required for AEP activation and Tau N368 cleavage in the LC of Tau P301S mice

To investigate whether DOPEGAL is implicated in Tau pathology and LC neuronal degeneration, we crossed Tau P301S transgenic mice with DBH knockout (DBH −/−) mice that lack NE completely, and thus cannot produce DOPEGAL, and found Tau hyperphosphorylation and N368 cleavage at different ages. DBH +/− littermates with normal NE levels were used as controls (33). At 3 months, AT8 and Tau N368 immunoreactivity in the LC was minimal in both Tau P301S/DBH +/− and Tau P301S/DBH −/− mice. While the abundance of pathological Tau escalated in both genotypes at 6-9 months, AT8, Tau N368, and Gallyas-Braak silver staining were significantly reduced in Tau P301S/DBH −/− mice (Figure 3A-C). Consistent with our finding that suppression of NE production protected against Tau toxicity *in vitro* (Supplemental Figure 3), TH staining revealed a loss of LC neurons in Tau P301S/DBH +/− mice at 9 months that was partially abrogated by DBH −/− (Figure 3B, D). To explore whether manipulation of NE synthesis affects cognitive impairment in Tau P301S mice, we tested spatial learning and memory using the Morris water maze (MWM). Comparable age-dependent deficits were observed at 3 and 6 months in NE-deficient and NE-competent Tau P301S mice. However, at 9 months, the performance of Tau P301S/DBH −/− mice was significantly better than Tau P301S/DBH +/− mice on several measures, including distance to platform during training and percentage of time spent in the target quadrant during a probe trial (Figure 3E-G). Similar results were observed with cued and contextual fear conditioning, which test hippocampal-independent and -dependent associative learning and memory, respectively (Figure 3H, I). AT8 and ThioS staining in hippocampus also demonstrated that Tau pathology was increased age-dependently in Tau P301S/DBH +/− mice compared to Tau P301S/DBH −/− mice (Supplemental Figure 5D). Thus, the full expression of Tau pathology and cognitive impairment in Tau P301S mice depends on NE.

**Figure 3.**
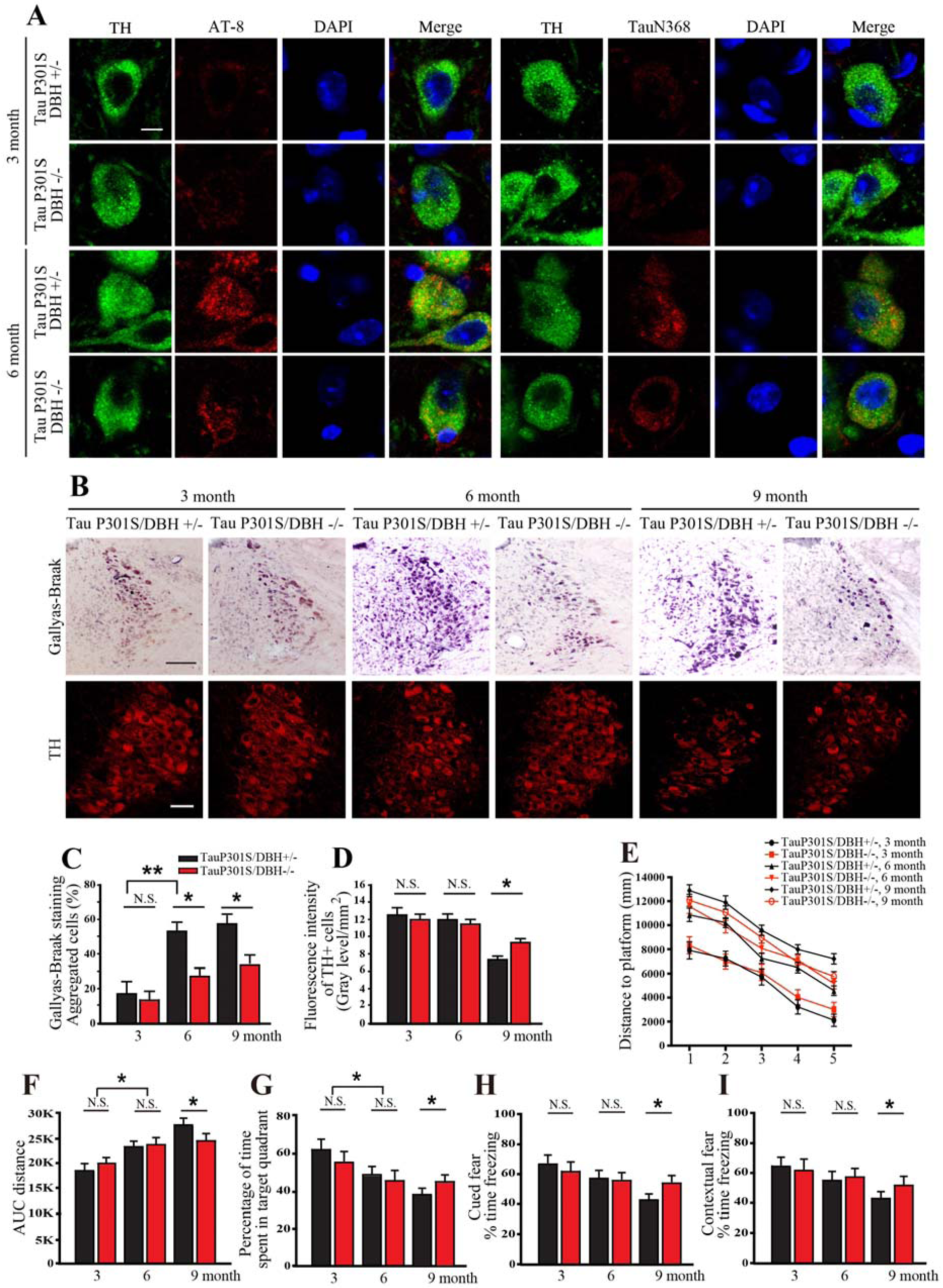
LC Tau pathology and cognitive impairment in P301S mice are dependent on NE. Tau P301S/DBH +/− and Tau P301S/DBH −/− mice at various ages were examined for Tau pathology in the LC by immunohistochemistry and cognitive impairment using Morris water maze and fear conditioning. **A**. Representative images of staining for tyrosine hydroxylase (TH; green), AT-8 or Tau N368 (red), and DAPI (blue). Scale bar = 20 μm. **B**. Representative Gallyas-Braak (upper panels) and TH staining. Scale bar = 100 μm. Quantification of **C.** Gallyas-Braak and **D.** TH staining in **B**. Data are shown as mean ± SEM. N=3 per group. * p<0.05. **E.** Distance travelled to the platform, **F.** area under the curve for total distance travelled, and **G.** percent time spent in the quadrant previously containing the platform during the probe trial in the Morris water maze, and percent time spent freezing during the **H.** cued fear and **I.** contextual fear tests following fear conditioning. All data were analyzed using one-way ANOVA and shown as mean ± SEM, N = 8 per group. * p < 0.05.

We next examined the impact of NE depletion on Tau pathology in the LC triggered by Tau cleavage. We reasoned that if DOPEGAL promotes the activation of AEP, then DBH −/− mice that cannot produce DOPEGAL would be protected from Tau pathology. We injected AAV encoding human Tau P301S or AEP-resistant Tau P301S N255A/N368A into the LC of DBH +/− and DBH −/− mice, and conducted IF co-staining with TH/AT8 and TH/Tau N368, as well as Gallyas-Braak silver staining. As expected, Tau P301S AAV elicited much greater Tau pathology than uncleavable Tau P301S N255A/368A virus. Strikingly, we found that the ability of Tau P301S to trigger AT8, N368, and Gallyas-Braak staining was attenuated significantly in DBH −/− mice (Supplemental Figure 5A-C). These data suggest that NE promotes Tau P301S N368 cleavage by AEP and its subsequentent phosphorylation and aggregation.

### DOPEGAL is toxic to LC neurons *in vivo*

MAO-A and DOPEGAL are upregulated in the LC of postmortem AD brains (24) and DOPEGAL, but not NE or its other oxidative or O-methylated metabolites, is toxic to differentiated catecholaminergic PC12 cells (23). To explore whether DOPEGAL is toxic to LC neurons *in vivo* and the potential involvement of AEP and Tau, we infused DOPEGAL (0.25 μg) into the LC of 2-month-old WT, Tau knockout (Tau −/−), MAPT transgenic mice that overexpress wild-type human Tau, and MAPT/AEP −/− mice. Four days following DOPEGAL administration, IF co-staining with TH and TUNEL revealed diminished TH and greater apoptosis in MAPT mice compared to WT, and these effects were abrogated by Tau or AEP depletion (Figure 4A-C). These results indicate that DOPEGAL-induced degeneration of LC neurons *in vivo* is partially mediated by a Tau- and AEP-dependent mechanism.

**Figure 4.**
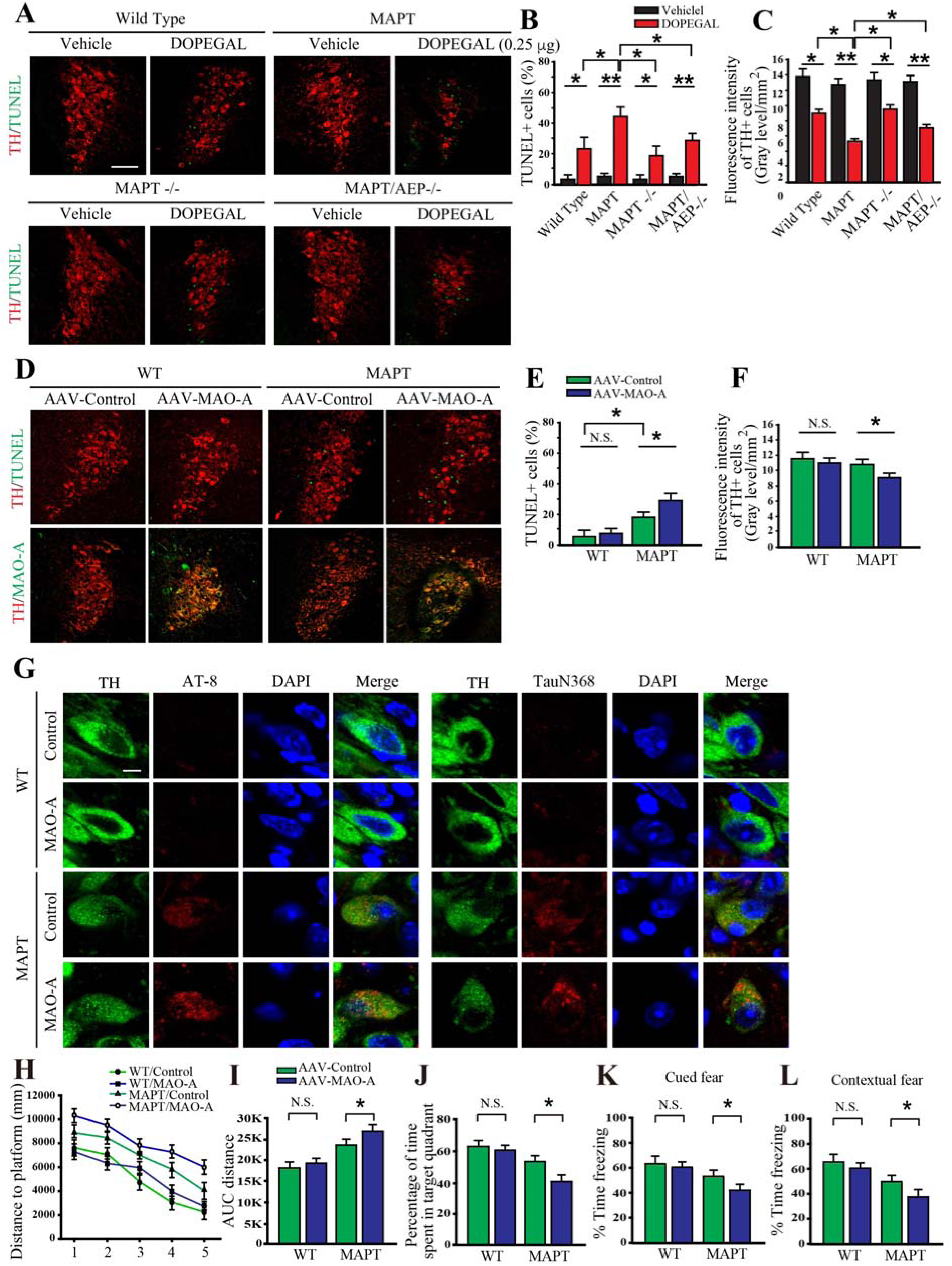
Increasing DOPEGAL in the LC induces Tau pathology, cell death, and cognitive impairment in MAPT transgenic mice. DOPEGAL (0.25 μg) was injected into the LC regions of wild-type, MAPT −/−, MAPT, or MAPT/AEP−/− mice. **A.** Representative images of TH (red) and TUNEL (green) staining in the LC 4 d following DOPEGAL injection. Scale bar = 100 μm. **B.** Quantification of TUNEL+ cells. **C.** Quantification of TH intensity. Data are shown as mean ± SEM. N=4 per group. * p<0.05, **p<0.01. AAV-MAO-A was injected into LC regions of wild-type or MAPT mice. **D.** Representative images of TH (red) and TUNEL (green) staining in the LC 3 months following AAV-MAO-A injection. Scale bar = 100 μm. **E.** Quantification of TUNEL. **F.** Quantification of TH intensity. Data are shown as mean ± SEM. N=6 per group. * p<0.05. **G**. Representative images of staining for TH (green), AT-8 or Tau N368 (red), and DAPI (blue). Scale bar = 20 μm. **H.** Distance travelled to the platform, **I.** area under the curve for total distance travelled, and **J.** percent time spent in the quadrant previously containing the platform during the probe trial in the Morris water maze, and percent time spent freezing during the **K.** cued fear and **L.** contextual fear tests following fear conditioning. All data were analyzed using one-way ANOVA and shown as mean ± SEM, N = 8 per group. * p < 0.05.

To investigate whether acceleration of endogenous NE metabolism and DOPEGAL production would have similar effects, we injected AAV-MAO-A or control virus into the LC of WT and MAPT mice and assessed pathology 3 months later. MAO-A overexpression failed to trigger neuronal loss in WT mice but significantly enhanced apoptosis and loss of TH-positive cells in MAPT mice (Figure 4D-F), which was accompanied by AT8 and Tau N368 staining in LC neurons (Figure 4G). The staining of AT8 and ThioS in hippocampus also demonstrated that Tau pathology was increased by MAO-A overexpression in MAPT mice compared to WT mice (Supplemental Figure 5E). MWM and fear conditioning tests demonstrated that MAO-A triggered significant cognitive dysfunction in MAPT mice compared to control virus, whereas it had no effect on WT mice (Figure 4H-L). These data suggest that Tau exacerbates DOPEGAL-mediated LC neuron toxicity and cognitive impairment.

### LC-derived human Tau can spread to interconnected brain regions in MAPT and 3xTg mice

Because the LC is the first brain structure to develop Tau lesions and has widespread connections to other areas of the brain, and Tau is capable of trans-synaptic propagation, LC neurons have been proposed to be the critical initiators of the stereotypical spread of Tau pathology in AD (4, 5). To explore this idea, we injected 3-month-old MAPT mice with either AAV-mCherry alone or AAV-mCherry + AAV-Tau under control of the noradrenergic-specific PRSx8 promoter into the LC (34), and then monitored Tau pathology throughout the brain 3 months later (Fig 5A). mCherry (red) and TH (blue) IF co-staining confirmed viral targeting to noradrenergic LC neurons, and AT8 (green) and N368 (green) Tau immunoreactivity were evident in the mice that received AAV-mCherry/AAV-Tau, but not mCherry alone (Figure 5B). AT8 immunohistochemistry (IHC) revealed that aberrant Tau had spread from the LC to the cerebellum, midbrain, hippocampus, entorhinal cortex and cortex (Figure 5A). Moreover, IF and Gallyas-Braak staining indicated that hyperphosphorylated Tau, AEP-cleaved Tau, and aggregated Tau were present in the entorhinal cortex (EC), hippocampus (HC) and cortex (Cx) of AAV-Tau injected mice (Figure 5C & D). By contrast, neither aberrant Tau nor mCherry were detected in the forebrain of mice injected with AAV-mCherry alone (Figure 5C), indicating that Tau pathology, rather than the virus itself, spread from the LC to these distal regions. Similar results were obtained when we performed the identical experiment in 3xTg mice (Supplemental Figure 6A-C).

**Figure 5.**
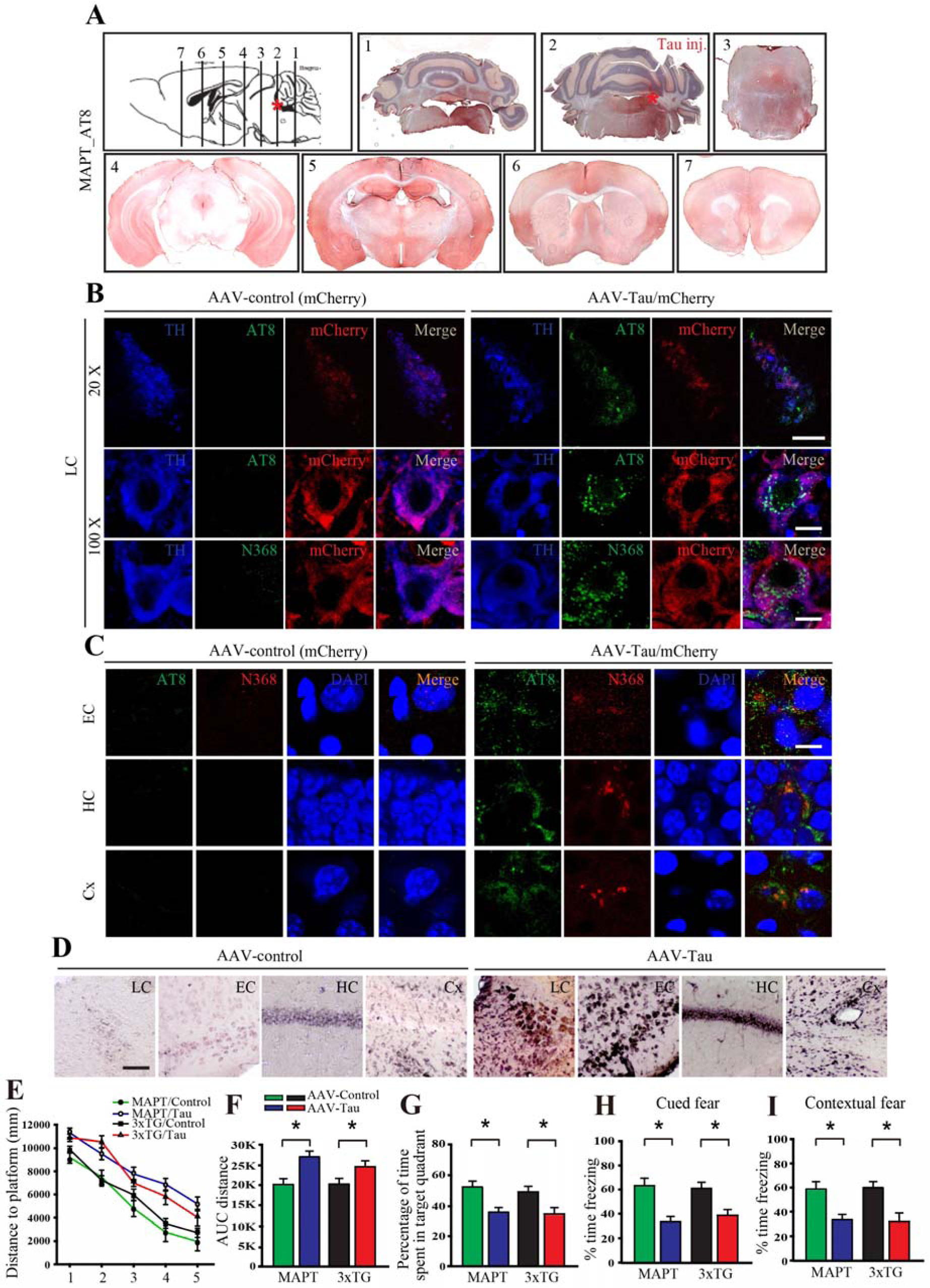
Viral-mediated Tau expression in the LC drives propagation of pathology to the forebrain in MAPT transgenic mice. LC-specific AAV-PRSx8-Tau + AAV-PRSx8-mCherry, or AAV-PRSx8-mCherry alone were injected into the LC of MAPT and 3xTG mice, and mice were assessed for Tau pathology throughout the brain by immunohistochemistry and cognitive impairment using Morris water maze and fear conditioning mice 3 months later. **A**. Representative images of AT8 immunostaining throughout the brain of MAPT mice. **B**. Representative images of immunofluorescent staining for TH (blue), AT8 or Tau N368 (green), and mCherry (red) in the LC of MAPT mice. Scale bar = 100 μm (top panels) and 20 μm (bottom panels). **C.** Representative images of immunofluorescent staining for AT8 (green) or Tau N368 (gred), and DAPI (blue) in the entorhinal cortex (EC), hippocampus (HC), and frontal cortex (Cx) of MAPT. Scale bar = 20 μm. **D**. Representative Gallyas-Braak staining in the LC, EC, HC, and Cx of MAPT mice. Scale bar = 100 μm. **E.** Distance travelled to the platform, **F.** area under the curve for total distance travelled, and **G.** percent time spent in the quadrant previously containing the platform during the probe trial in the Morris water maze, and percent time spent freezing during the **H.** cued fear and **I.** contextual fear tests following fear conditioning in MAPT and 3xTg mice. All data were analyzed using one-way ANOVA and shown as mean ± SEM, N = 8 per group. * p < 0.05 compared with control group.

To test whether pathogenic Tau originating in the LC can induce cognitive deficits, we conducted MWM and fear conditioning 3 months after intra-LC Tau virus administration in MAPT and 3xTg mice. Tau-injected mice spent significantly less time in the target quadrant during the MWM probe trial (Figure 5E-G) and were impaired in both cued and contextual freezing following fear conditioning (Figure 5H-I). Together, these behavioral tests indicate that LC-derived Tau pathology can spread to the forebrain and produce cognitive impairment. AEP cleavage of Tau increases its aggregation and neurotoxicity, and depletion of AEP diminishes Tau pathology in Tau P301S mice (18). Tau cleavage by AEP is necessary and sufficient for its neurotoxicity in LC neurons (Supplemental Figure 2), and the appearance of AEP-cleaved Tau N368 is tightly associated with AT8 immunoreactivity, suggesting that AEP cleavage of Tau may facilitate its spread. We prepared primary LC neurons from neonatal WT and AEP −/− mice and infected them with AAV-PRSx8-Tau or control virus. IF co-staining with TH and Tau5 or the human-specific Tau HT7 antibody showed that human Tau was specifically overexpressed in the LC neurons (Supplemental Figure 7A, upper panels). TH and N368 or AT8 co-staining demonstrated that human Tau was truncated at N368 and hyperphosphorylated in WT LC neurons, whereas both N368 and AT8 immunoreactivity were attenuated in AEP −/− LC neurons (Supplemental Figure 7A, lower panels). To determine whether AEP is necessary for Tau aggregation and spread *in vivo*, we injected 3-month-old WT and AEP −/− mice with AAV-PRSx8-Tau + AAV-PRSx8-mCherry in the LC and monitored Tau pathology in the LC and forebrain 3 months later. AT8, N368 Tau, and Gallyas-Braak staining indicated that hyperphosphorylated and AEP-cleaved Tau accumulation in the LC and subsequent spread to the HC, Cx, and EC were largely retarded in AEP −/− mice (Figure 6A-C). A schematic map of Tau pathology spread from the LC in MAPT, 3xTg, WT, and AEP −/− mice as viewed by AT8 staining is shown in Supplemental Figure 7B. Moreover, deletion of AEP prevented cognitive impairment induced by Tau overexpression in the LC (Figure 6D-H). These data indicate that AEP contributes to the spread of LC-derived Tau pathology to the forebrain and associated cognitive deficits.

**Figure 6.**
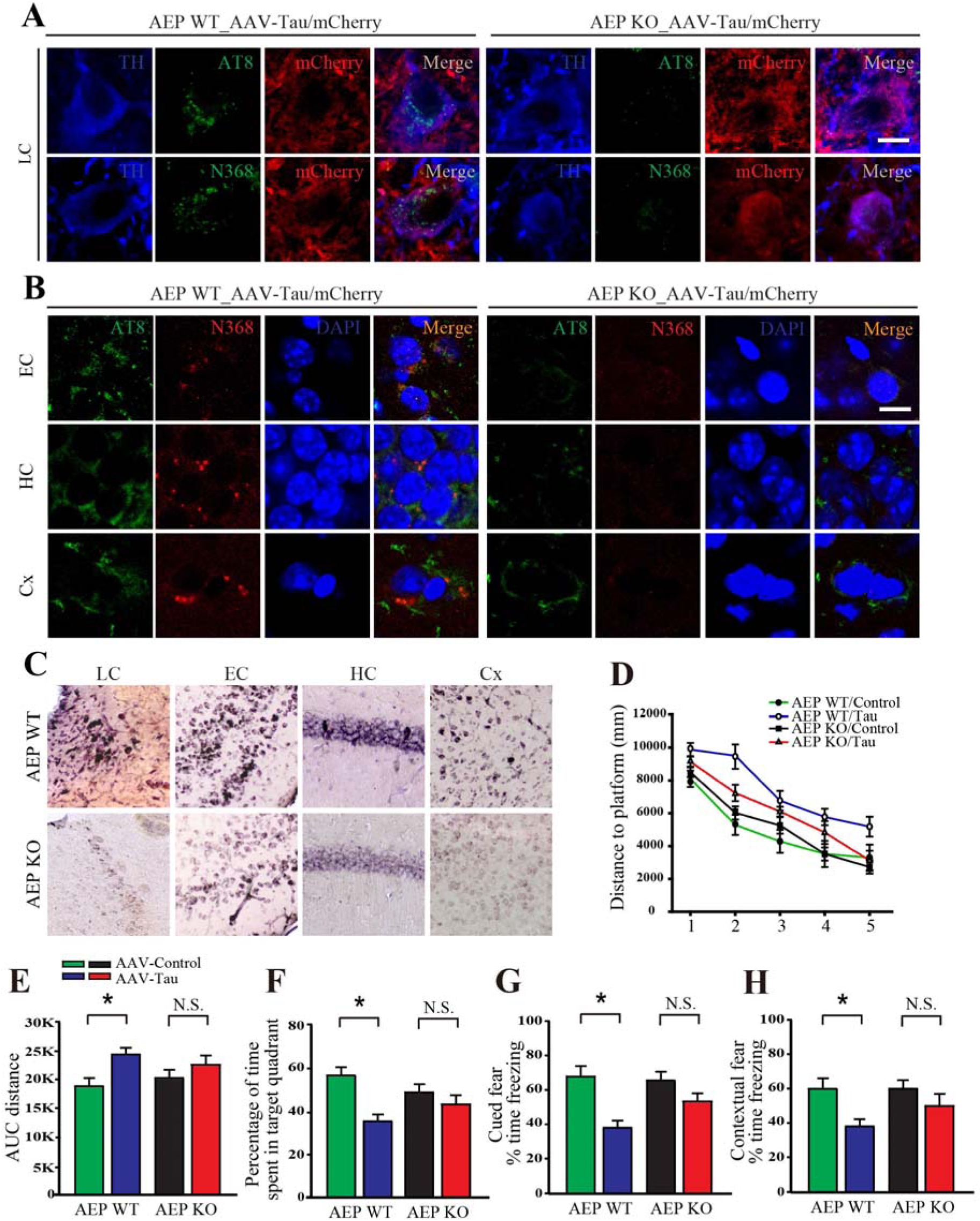
Viral vector-mediated Tau pathology in the LC and its propagation to the forebrain are blunted in AEP −/− mice. LC-specific AAV-PRSx8-Tau + AAV-PRSx8-mCherry, or AAV-PRSx8-mCherry alone were injected into the LC of AEP +/+ or AEP −/− mice, and pathology and cognition were assessed 3 months later. **A**. Representative images of immunofluorescent staining of tyrosine hydroxylase (TH, blue), AT8 or N368 (green), and mCherry (red) in the LC. Scale bar = 20 μm. **B**. Representative images of immunofluorescent staining of AT8 (green), N368 (red), and DAPI (blue) in the entorhinal cortex (EC), hippocampus (HC), and cortex (Cx). Scale bar = 20 μm. **C**. Representative Gallyas-Braak staining in the LC, EC, HC, and Cx of AEP +/+ mice. Scale bar = 100 μm. **D.** Distance travelled to the platform, **E.** area under the curve for total distance travelled, and **F.** percent time spent in the quadrant previously containing the platform during the probe trial in the Morris water maze, and percent time spent freezing during the **G.** cued fear and **H.** contextual fear tests following fear conditioning. All data were analyzed using one-way ANOVA and shown as mean ± SEM. N = 8 per group. * p < 0.05 compared with control group.

## Discussion

Pathogenic forms of Tau first appear in noradrenergic LC neurons (3–6), which degenerate later in AD, and LC dysfunction contributes to AD-like neuropathology and cognitive deficits (6, 8, 9, 11–13, 16). However, the mechanism underlying the selective vulnerability of LC neurons has remained elusive. In the current study, we report that DOPEGAL, the NE metabolite produced by MAO-A oxidation, triggers AEP activation via oxidative stress, resulting in Tau N368 cleavage that is prone to hyperphosphorylation, aggregation, and propagation, and triggers noradrenergic neuronal death. In addition, DOPEGAL, but not NE, facilitates the fibrilization of purified recombinant Tau. Tau hyperphosphorylation and N368 cleavage were age-dependently increased in the LC of both MAPT and 3xTg mice, consistent with our previous finding that AEP is upregulated and activated in the brain during aging. Importantly, we made the same observations in the LC neurons of human AD brains, indicating that active AEP contributes to Tau pathology in the LC.

DOPEGAL is an appealing candidate to explain the selective vulnerability of LC neurons in AD for several reasons. First, it is only produced in noradrenergic cells. Second, it is increased in the LC of human AD brains and is toxic to noradrenergic neurons. Finally, the close neurochemical relationship between NE and DA suggests that the toxicity of the DA MAO metabolite DOPAL may have a correlate in the noradrenergic system. Previous studies show that physiological concentrations of DOPAL trigger the formation of α-Syn oligomers and aggregates in both a cell-free system and in cell culture, and produce cytotoxicity *in vitro* and *in vivo*. We recently reported that DOPAL activates AEP in dopaminergic neurons and leads to α-synuclein N103 cleavage by AEP, resulting in its aggregation and dopaminergic neuronal degeneration in the SN (21). Thus, DOPAL-α-Syn interactions may underlie the selective vulnerability of DA neurons in Parkinson’s disease (35–39), and we suspected that a similar interaction might occur between DOPEGAL and Tau in AD. Here we present multiple converging lines of evidence implicating DOPEGAL and AEP in the vulnerability of LC neurons to Tau pathology and toxicity. First, DOPEGAL, but not NE itself, increased AEP activity and cell death. Second, overexpression of MAO-A, which oxidizes NE into DOPEGAL, facilitated AEP activity, Tau N368 cleavage, and cell death, while decreasing MAO activity or blocking Tau cleavage had the opposite effect. Third, preventing DOPEGAL production via DBH depletion or Tau cleavage by AEP −/− mitigated the deleterious effects of Tau on LC neurons, whereas preventing the breakdown of DOPEGAL by AR or ALDH inhibition escalated AEP activity and cell death. Finally, full DOPEGAL toxicity was dependent on the presence of Tau. Together, these data suggest that DOPEGAL stimulates AEP activation via oxidative stress, resulting in Tau N368 cleavage and aggregation, LC neuronal degeneration, and the spread of Tau pathology to the forebrain and associated cognitive impairment. A schematic model is proposed in Figure 7. Given that DOPAL can directly modify α-Syn, it is possible that DOPEGAL may covalaently modify the lysine residues and oxidize Met groups on Tau, thus facilitating its conformational change and fibrillization. We did find that DOPEGAL accelerated the aggregation of recombinant Tau *in vitro*, but further experiments will be required to pinpoint the covalent and/or non-covalent mechanisms.

**Figure 7.**
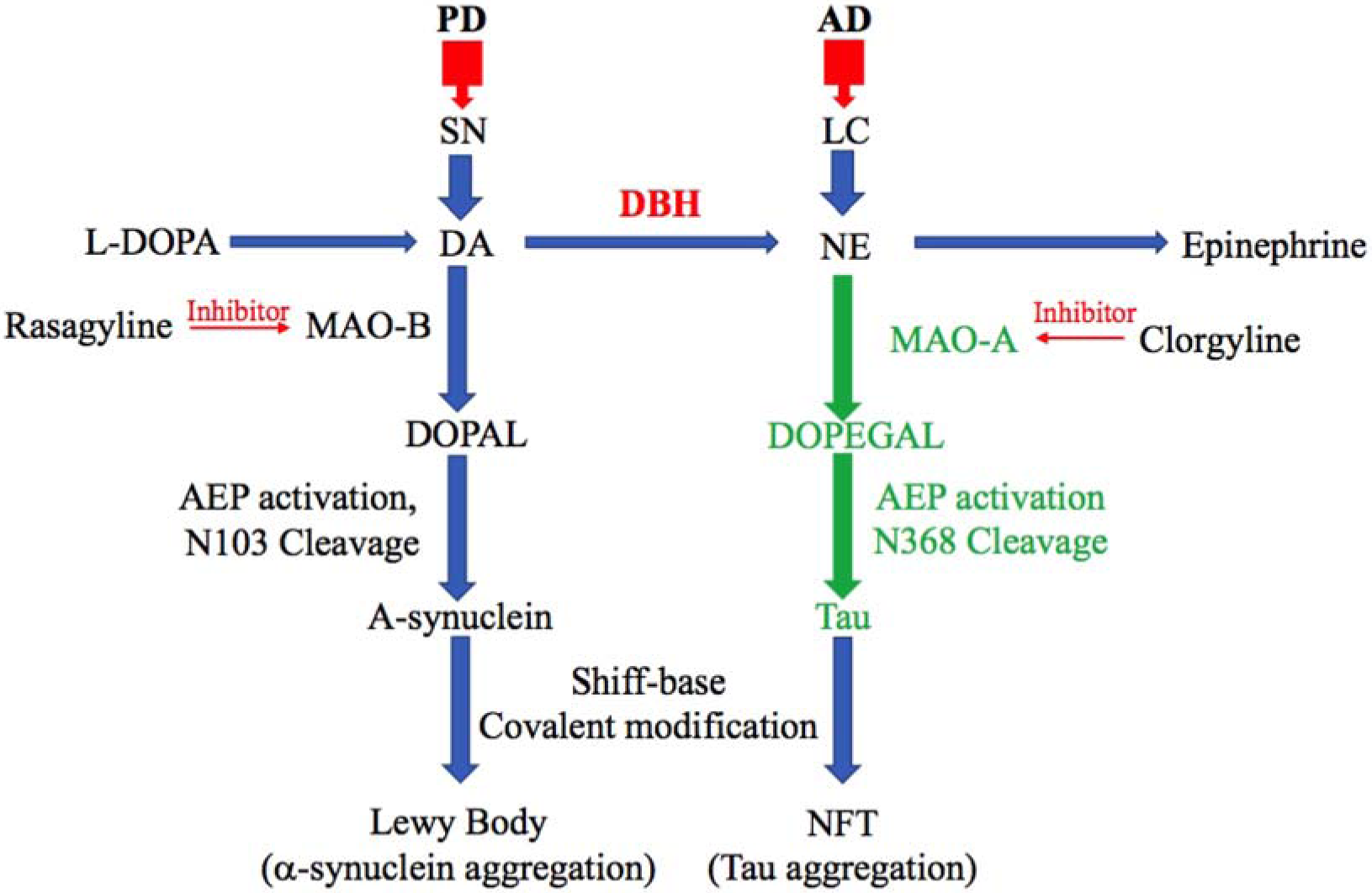
A Schematic model for the selective vulnerability by DA or NE metabolites, monoamine oxidase, and AEP in Parkinson’s disease and Alzheimer’s disease. DA metabolite by MAO-B, DOPAL, have the toxic effect in PD pathology specifically in substantia nigra, leading to AEP activation and alpha-synuclein toxicity. Similarly, MAO-A and its toxic metabolite of NE, DOPEGAL, induce tau toxicity in noradrenergic cells of LC in the early stage of AD progression.

MAO-A preferentially oxidizes serotonin (5-HT) and NE, whereas MAO-B preferentially oxidizes phenylethylamine (PEA), and both forms can oxidize DA (40). Increases of both MAO-A and MAO-B have been detected in human AD brains. For instance, DOPEGAL and MAO-A levels are elevated 2.8- and 3.6-fold in AD LC neuronal cell bodies compared to controls (24), while MAO-B expression and activity are higher in multiple brain regions and cell types in AD (41, 42). While both MAO-A and MAO-B were capable of increasing AEP activity, Tau N368 cleavage, and cell death in our experiments, we favor MAO-A as the main driver of DOPEGAL production *in vivo*, with MAO-B playing a greater role generating the neurotoxic DA metabolite DOPAL in PD (21, 43) (Figure 7); that said, confirming this hypothesis will require additional experiments.

Beause DOPEGAL, but not NE itself, generates free radicals and activates mitochondrial permeability transition (27), DOPEGAL may contribute to the loss of noradrenergic LC neurons in AD (44). Employing Tau −/− neurons and mice, we showed that DOPEGAL-induced neurotoxicity is partially dependent on Tau. Knockout of AEP significantly ameliorated the LC neuronal degeneration and Tau pathology elicited by DOPEGAL exposure in MAPT mice, indicating that AEP cleavage of Tau is an important factor in LC vulnerability. This finding is further supported by our experiments with the AEP cleavage-resistant Tau P301S/N255/368A mutant. In SH-SY5Y cells, we found that the ability of AEP-cleaved Tau N368 to trigger cell death was more robust than WT Tau and comparable to P301S Tau. By contrast, introducing the N255/368A mutations onto a P301S background substantially reduced the toxicity of P301S Tau. Moreover, P301S Tau strongly induced Tau pathology in LC neurons *in vivo* that was associated with Tau N368 cleavage, while AEP-resistant P301S/N255/368A Tau was much less active. P301S-induced Tau pathology was also attenuated in the LC of DBH −/− mice, indicating that NE is required for Tau to exert its full neurotoxicity. This interpretation is further supported by our findings that siRNA-mediated knockdown of DBH (and thus NE) production blunted Tau-induced cell loss in catecholaminergic SH-SY5Y cells. Combined, these data support a model whereby LC vulnerability in AD is caused by NE metabolism to DOPEGAL by MAO-A, which activates AEP and Tau cleavage, leading to Tau pathology and neuronal death. Previous research has shown that Tau pathology is detectable in 3xTg mice at 9-12 months (45) and in MAPT mice at 12 months (46), so in our experiments, any appearance of aberrant Tau at 6 months must be derived from the LC. We observed the appearance of hyperphosphorylated, AEP-cleaved, and Gallyas-Braak-positive forms of Tau in the EC, HC, and Cx of both transgenic strains following intra-LC infusion of mCherry + Tau virus. Importantly, mCherry did not spread to distal brain regions, and there was no forebrain Tau pathology detected in mice that received mCherry alone. These results clearly show that, at least in these models, Tau pathology originating in the LC is capable of propagating to interconnected brain regions, including those that also show early vulnerability in AD. To determine whether this spread of pathogenic Tau had any functional effects, we tested mice for signs of cognitive impairment and found that the performances of 3xTg and MAPT mice in the Morris water maze and fear conditioning assays worsened following Tau expression in the LC and subsequent spread to the forebrain. Remakably, both the spread of Tau pathology and the accelerated cognitive deficits following Tau expression in the LC were alleviated in AEP −/− mice. Here we provide a combination of *in vitro*, cellular, *in vivo*, genetic, and pharmacological evidence to describe a novel neurochemical/molecular pathway involving NE metabolism and AEP that may explain the selective vulnerability of LC neurons to developing Tau pathology, the later demise of these cells, and the potential spread of pathology from the LC to the forebrain. Identification of the DOPEGAL-AEP-Tau N368 pathway lays the foundation for development of LC/AEP-based therapies that could slow the progression of AD.

## Methods

### Animals

Tau P301S mice on a C57BL/6J background (line PS19), MAPT mice, 3XTg mice, and wild-type C57BL/6J mice were obtained from the Jackson Laboratory (stock numbers 008169, 005491, 34830, and 000664, respectively). The AEP −/− mice were manintained on a mixed 129/Ola and C57BL/6 background. DBH −/− mice were maintained on a mixed 129/SvEv and C57BL/6 background. Animal care and handling was performed according to the Declaration of Helsinki and Emory Medical School guidelines. Tau P301S mice were crossed with DBH −/− mice to generate Tau P301S/DBH −/− mice. Adrenergic agonists and/or L-DOPS were used to rescue the DBH −/− embryonic lethality and male mating deficit, as described (47). MAPT mice were crossed with AEP −/− mice to generate MAPT/AEP −/− mice. Investigators were blinded to the group allocation during the animal experiments. The protocol was reviewed and approved by the Emory Institutional Animal Care and Use Committee.

### Human tissue samples

Paraffin sections containing the LC from 5 postmortem AD cases (age 74.5 ± 11.2 years, mean ± SD) and 5 cognitively normal controls (age 73.9 ± 12.7 years) were obtained from the Emory Alzheimer’s Disease Research Center. The study was approved by the Biospecimen Committee. AD was diagnosed according to the criteria of the Consortium to Establish a Registry for AD and the National Institute on Aging. Diagnoses were confirmed by the presence of amyloid plaques and neurofibrillary tangles in formalin-fixed tissue. Informed consent was obtained from the subjects prior to their deaths.

### Transfection and viral infection of cells

HEK293 cells and SH-SY5Y cells were transfected with plasmids encoding Tau, SiDBH, MAO-A, or MAO-B using polyethylenimine (PEI) or Lipofectamine 3000 (Thermo Fisher Scientific). AAV-Tau, AAV-Tau P301S, AAV-Tau 1-368, and AAV Tau N225/368A were used for infection in SH-SY5Y cells. Neurotoxicity was analyzed using LDH assay (CytoTox 96® Non-Radioactive Cytotoxicity Assay, Promega).

### Western blot analysis

Cultured cells were lysed in lysis buffer (50 mM Tris, pH 7.4, 40 mM NaCl, 1 mM EDTA, 0.5% Triton X-100, 1.5 mM Na_3_VO_4_, 50 mM NaF, 10 mM sodium pyrophosphate, 10 mM sodium β-glycerophosphate, supplemented with a cocktail of protease inhibitors), and centrifuged for 15 min at 16,000 g. The supernatant was boiled in SDS loading buffer. After SDS-PAGE, the samples were transferred to a nitrocellulose membrane. Primary antibodies to the following targets were used: Tau5, AT-8, HT7 (Invitrogen), beta-actin (Sigma-Aldrich), MAO-B (GeneTex), MAO-A (GE healthcare), TH (Sigma-Aldrich), AEP, and Tau N368 (made in-house).

### AEP activity assay

Assay buffer (20 mM citric acid, 60 mM Na_2_HPO_4_, 1 mM EDTA, 0.1% CHAPS and 1 mM DTT, pH 6.0) of 100 μl including 100 μM AEP substrate Z-Ala-Ala-Asn-AMC (ABchem) was added to cell or tissue lysates. AMC released by substrate cleavage was quantified using a fluorescence plate reader at 460 nm for 1 h in kinetic mode.

### MAO-A activity assay

Cell or tissue lysates (20 μg) were incubated with the working solution of 100 μl containing 400 μM Amplex® Red reagent, 2 U/ml HRP, and 2 mM p-tyramine substrate with MAO-B inhibitor, pargyline (Molecular Probes). The fluorescence of MAO-A activity was measured in a fluorescence plate reader using excitation in the range of 530-560 nm and emission at 590 ±10 nm at 37°C for 2 h in kinetic mode.

### Thioflavin T (ThioT) assay

ThioT stock solution (3 mM) was filtered through a 0.2-μm syringe filter (Sigma-Aldrich. Cat#T3516). The stock solution was diluted into the aggregation buffer (20 mM Tris, pH7.4, 100 mM NaCl, 1 mM EDTA) to generate the working solution (30 μM). ThioT working solution with heparin (30 μM), recombinant Tau (7 µM), and DOPEGAL (500 μM) was incubated on an orbital shaker at 37°C and excited at 440 nm and emitted at 482 nm to measure the fluorescence intensity on the plater reader (BioTek, #251639, Vermont, USA). Fluorescence values (excitation at 440 nm and emission 482 nm) onto the plate reader (BioTek, #251639, Vermont, USA) were recorded for 5 d.

### Immunostaining

Paraffin-embedded human brain sections or free-floating mouse brain sections sliced by cryotome were treated with 0.3% H_2_O_2_ for 10 min. Sections were washed three times in PBS and blocked in 1% BSA, 0.3% Triton X-100, for 30 min, followed by overnight incubation with TH antibody (1: 1000), anti-Tau5, anti-Tau N368, or AT-8 antibody (1: 500) at 4°C. The signal was developed using a Histostain-SP kit (Invitrogen). For IF, sections were incubated overnight with various primary antibodies at 4°C. Then, the sections were incubated with the matched fluoroconjugated secondary antibody for 2 h at room temperature, followed by three washes in PBS. The slides were washed three times in PBS and covered with a glass using mounting solution, after DAPI staining for 5 min.

### Stereotaxic injection

AAV-PRSx8-mCherry, AAV-PRSx8-Tau, AAV-Tau P301S, AAV-Tau N255/368A, or AAV-MAO-A were injected into the LC region of mice. 3-month-old mice of each group were anesthetized with isoflurane (Piramal Healthcare). Meloxicam (2 mg/kg) was injected subcutaneously for analgesia (Loxicom, Norbrook). Unilateral intracerebral injection of virus was performed stereotaxically at the following coordinates: anteroposterior (AP) −5.4 mm and mediolateral (ML) −1.2 mm relative to bregma, and dorsoventral (DV) −3.7 mm from the dural surface. 2 μl of viral suspension was injected into each site using a 10-μl Hamilton syringe with a fixed needle at a rate of 0.25 μl/min. The needle remained in place for 5 min after the viral suspension was completely injected, and then was slowly removed over 2 min. The mice were placed on a heating pad until recovery from the anesthesia.

### Behavioral testing

#### Morris water maze

Mice were trained in a round, water-filled tub (52” diameter) in an environment with extra maze cues. Each subject was given 4 trials/d for 5 consecutive d, with a 15-min intertrial interval. The maximum trial length was 60 s, and if subjects did not reach the platform in the allotted time, they were manually guided to it. Following the 5 d of task acquisition, a probe trial was presented, during which time the platform was removed and the percentage of time spent in the quadrant that previously contained the escape platform during task acquisition was measured over 60 s. All trials were analysed for latency and swim speed by MazeScan (CleverSys, Inc.).

#### Fear conditioning

The ability to form and retain an association between an aversive experience and environmental cues was tested with a standard fear conditioning paradigm occurring over a period of 3 d. Mice were placed in the fear conditioning apparatus (7” W, 7” D X 12” H, Coulbourn Instruments), composed of Plexiglas with a metal shock grid floor, and allowed to explore the enclosure for 3 min. Following this habituation period, 3 conditioned stimulus (CS)-unconditioned stimulus (US) pairings were presented with a 1-min intertrial interval. The CS consisted of a 20-s 85-db tone, and the US consistedd of 2 s of a 0.5-mA footshock, which was co-terminated with each CS presentation. One min following the last CS-US presentation, mice were returned to their home cage. On day 2, the mice were presented with a context test, during which subjects were placed in the same chamber used during conditioning on Day 1, and the amount of freezing was recorded via camera and the software provided by Colbourn. No shocks were given during the context test. On day 3, a tone test was presented, during which time subjects were exposed to the CS in a novel compartment. Initially, animals were allowed to explore the novel context for 2 min. Then the 85-db tone was presented for 6 min, and the amount of freezing behavior was recorded.

### Quantification and statistical analysis

All cell culture data are expressed as mean ± S.E.M. from 3 or more independent experiments, and the level of significance between 2 groups was assessed with Student’s t-test. For experiments consisting of more than 2 groups, one-way ANOVA followed by LSD post hoc test was applied. A value of p < 0.05 was considered to be statistically significant. Sample sizes were determined by Power and Precision (Biostat).

## AUTHOR CONTRIBUTIONS

K.Y. conceived the project, designed the experiments, analyzed the data, and wrote the manuscript. S.S.K. designed and performed most of the experiments. X.L. prepared primary neurons and assessed with animal experiments. C.J. performed the crosses, pharmacological embryonic lethality rescue, and genotyping for experiments with the DBH −/− mice. E.H.A. and J.X. performed Tau purification and some of the *in vitro* experiments. F.P.M. prepared the uncleavable Tau virus. H.R. L., X. Y., and S.S.K. assisted with data analysis and interpretation and critically read the manuscript. D.W. helped design the experiments and edit the manuscript.

## ACKNOWLEDGMENTS

This work was supported by the NIH (R01AG051538 to K. Y. and RF1AG047667 to D.W.). We thank the Emory Alzheimer’s Disease Research Center for postmortem human AD and healthy control samples. We thank the University of Washington Alzheimer’s Disease Research Center (NIH P50AG005136) and the Kaiser Permanente Adult Changes in Thought Study (NIH U01 AG006781) research subjects and their families for brain donation, a tremendous gift to science. We thank Ms. Allison Beller for coordination of tissue requests, subject selection, and tissue transfer, Ms. Kim Howard for histological support, and Ms. Cheryl Strauss for helpful editing of the manuscript. This study was supported in part by the Rodent Behavioral Core (RBC) and Viral Vector Core, which are subsidized by the Emory University School of Medicine and are part of the Emory Integrated Core Facilities. Additional support was provided by the Emory Neuroscience NINDS Core Facilities (P30NS055077). Further support was provided by the Georgia Clinical & Translational Science Alliance of the National Institutes of Health under Award Number UL1TR002378.

## COMPETING FINANCIAL INTERESTRS

The authors have declared that no conflict of interests exists.

## Supplemental information

**Supplemental Figure 1.**
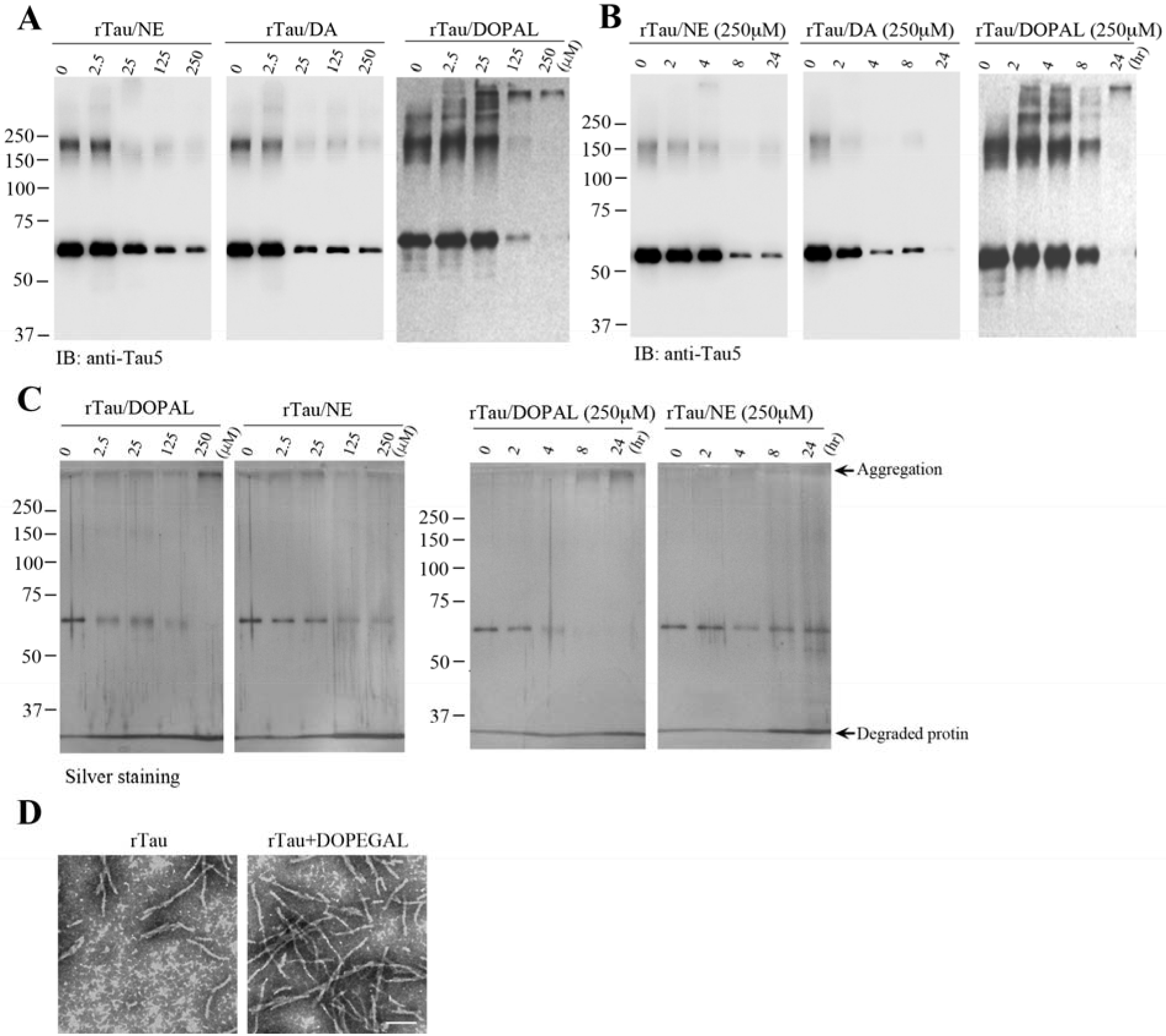
NE, DA, and DOPAL trigger Tau degradation *in vitro*. **A.** Recombinant Tau (1 μg) was incubated with NE, DA, or DOPAL of 0, 2.5, 25, 125, or 250 μM in a 37 °C shaker for 24 h. Immunoblotting showed that NE, DA, and DOPAL dose-dependently induced Tau degradation. **B.** NE, DA, and DOPAL time-dependently stimulated Tau degradation from 2 to 24 h. **C.** Silver staining confirmed that Tau was dose- and time-dependently degraded by NE and DOPAL. **D.** Enhanced Tau fabrilization by DOPEGAL was demonstrated by electron microscopy. Scale bar = 100 nm.

**Supplemental Figure 2.**
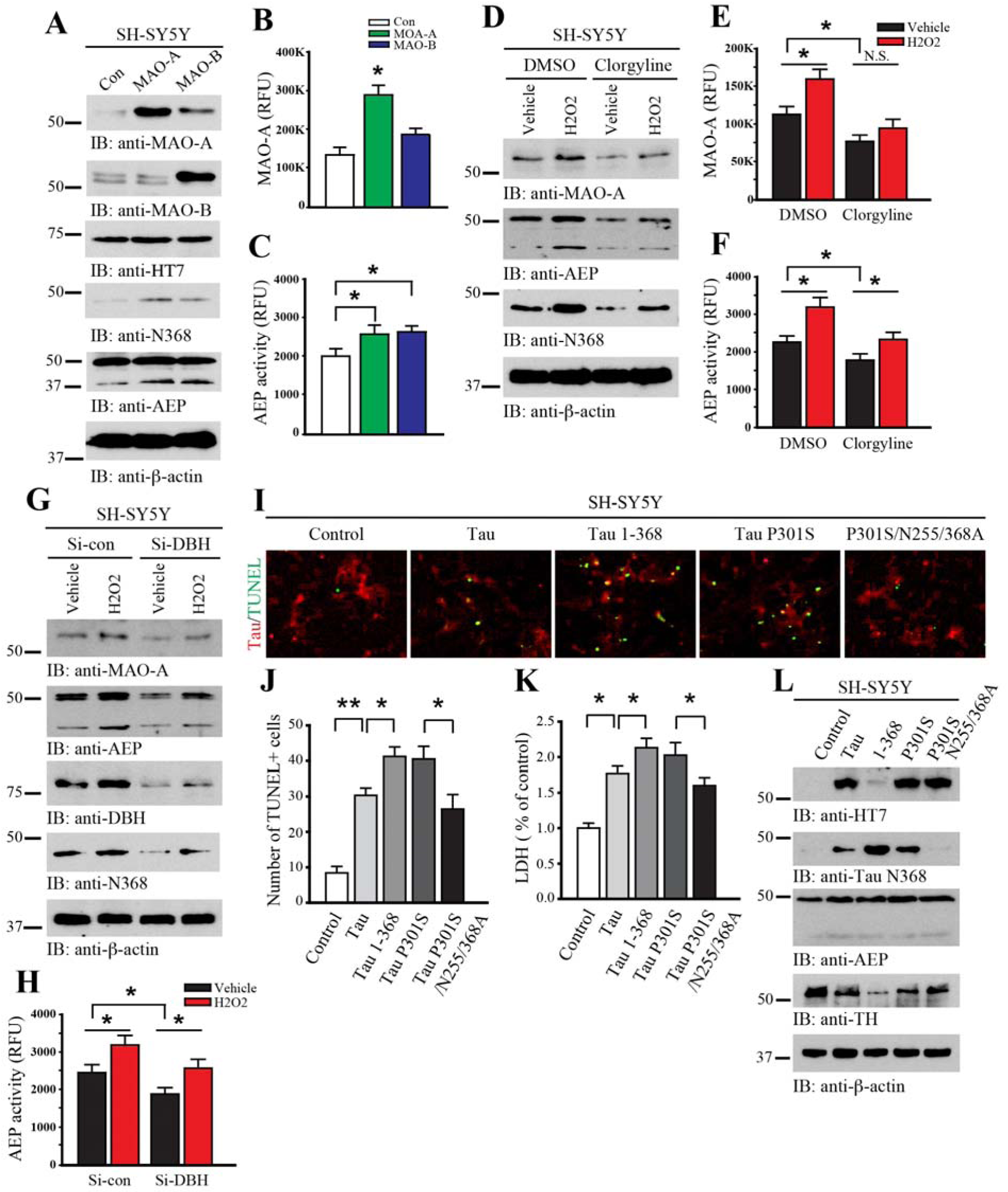
MAO-A or aberrant Tau overexpression induces AEP activation, Tau N368 cleavage, and cell death in SH-SY5Y cells. SH-SY5Y cells were transfected with MAO-A or MAO-B. **A.** Western blot analysis was conducted on cell lysates using antibodies against AEP and different forms of Tau with β-actin as a control. **B**. The activation of MAO-A in transfected cells was confirmed by enzymatic assay. **C**. AEP enzymatic assay showed that the overexpression of MAO-A and MAO-B activated AEP. Data are shown as mean ± SEM. N=3 per group. * p<0.05. H_2_O_2_ triggered MAO-A and AEP activation. SH-SY5Y cells were pretreated with Clorgyline (10 μM), followed by treatment with H_2_O_2_ (100 μM) for 4 h. **D**. Western blot analysis showed that MAO-A and AEP were enhanced by H_2_O_2_, and the effects of H_2_O_2_ were attenuated by the specific MAO-A inhibitor clorgyline. **E**. Activation of MAO-A by H_2_O_2_ was confirmed by MAO-A enzymatic assay. **F**. AEP enzymatic assay showed that H_2_O_2_ activation of AEP was inhibited by clorgyline. Data are shown as mean ± SEM. N=3 each group. * p<0.05. Inhibition of NE synthesis reduced H_2_O_2_-induced AEP activation. SH-SY5Y cells were transfected with siRNA for DBH or a control sequence, followed by treatment with H_2_O_2_ (100 μM) for 4 h. **G**. Western blot anaylsis showed that inhibition of NE synthesis by knocking down DBH reduced H_2_O_2_-induced AEP activation and Tau N368 cleavage. **H**. H_2_O_2_-induced AEP activity was inhibited by knocking down DBH. Data are shown as mean ± SEM. N=3 each group. * p<0.05. Tau cleavage by AEP induces SH-SY5Y cell death. SH-SY5Y cells were infected by AAV expressing Tau, Tau N368, Tau P301S or AEP-resistant Tau P301S/N255/368A. **I.** Representative images of cells co-stained for Tau (red) and TUNEL (green). **J & K**. Quantification of TUNEL+ cells and LDH assay showed that Tau-induced cell death was inhibited by AEP-resistant uncleavable Tau. Data are shown as mean ± SEM. N=3 per group. * p<0.05. **L**. Western blot analysis showed that TH cell loss was inhibited by uncleavable Tau P301S/N255/368A.

**Supplemental Figure 3.**
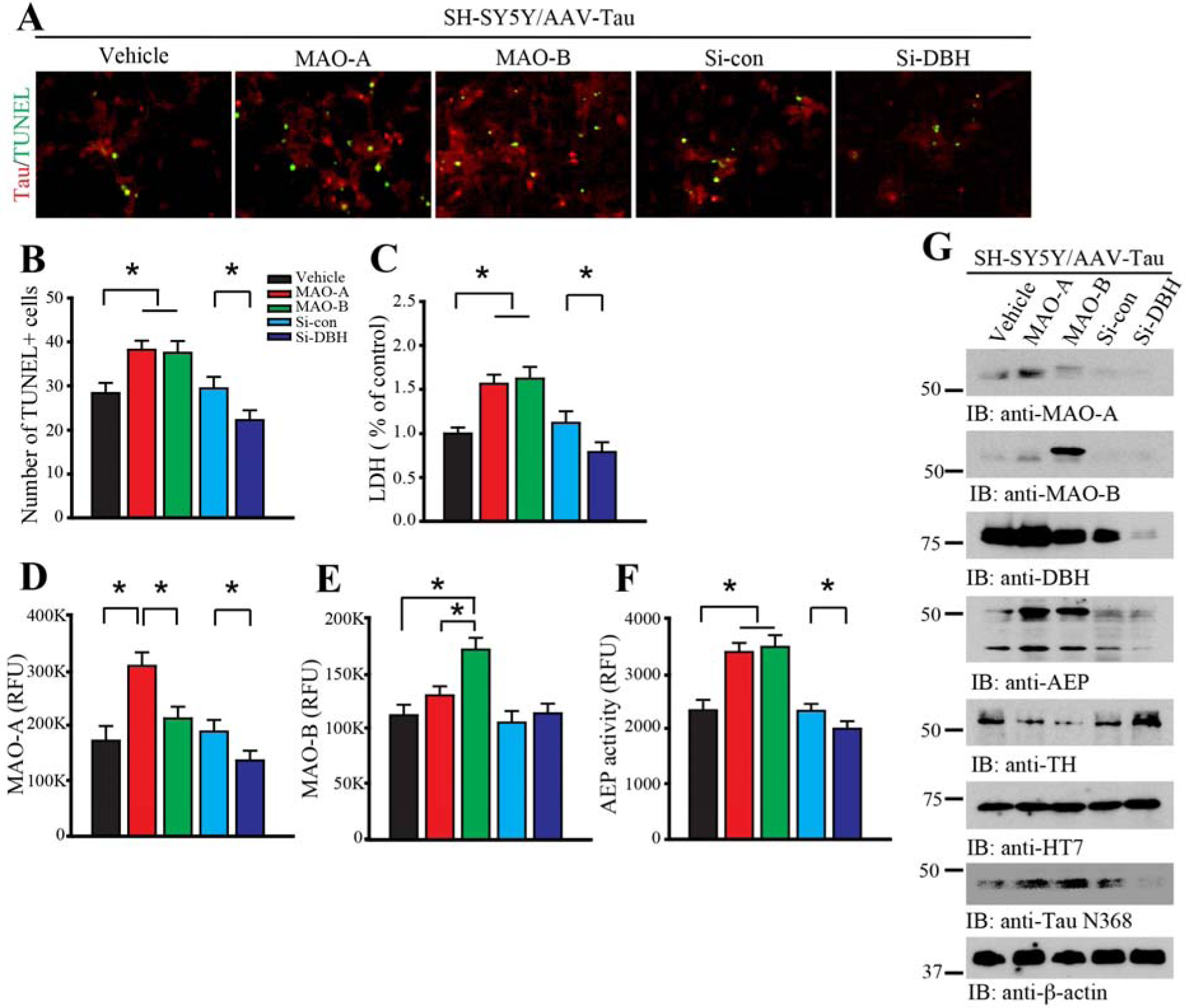
Tau-induced neurotoxicity in SH-SY5Y cells is enhanced by MAO overexpression and attenuated by DBH depletion. **A.** SH-SY5Y cells were infected by AAV expressing Tau with MAO-A or MAO-B, or transfected with DBH siRNA. TUNEL was co-stained with Tau. **B & C.** Quantification of TUNEL+ cells and LDH assay show that Tau-induced cell death is increased by MAO-A or MAO-B, and is reduced by knockdown of DBH. The activity change of MAO-A (**D**), MAO-B (**E**), and AEP (**F**) by AAV-MAO-A or siDBH was verified by enzymatic assay. Data are shown as mean ± SEM. N=3 per group. * p<0.05. **G**. Western blot analysis shows that MAO-A overexpression induces AEP activation, Tau cleavage, and TH reduction in SH-SY5Y cells.

**Supplemental Figure 4.**
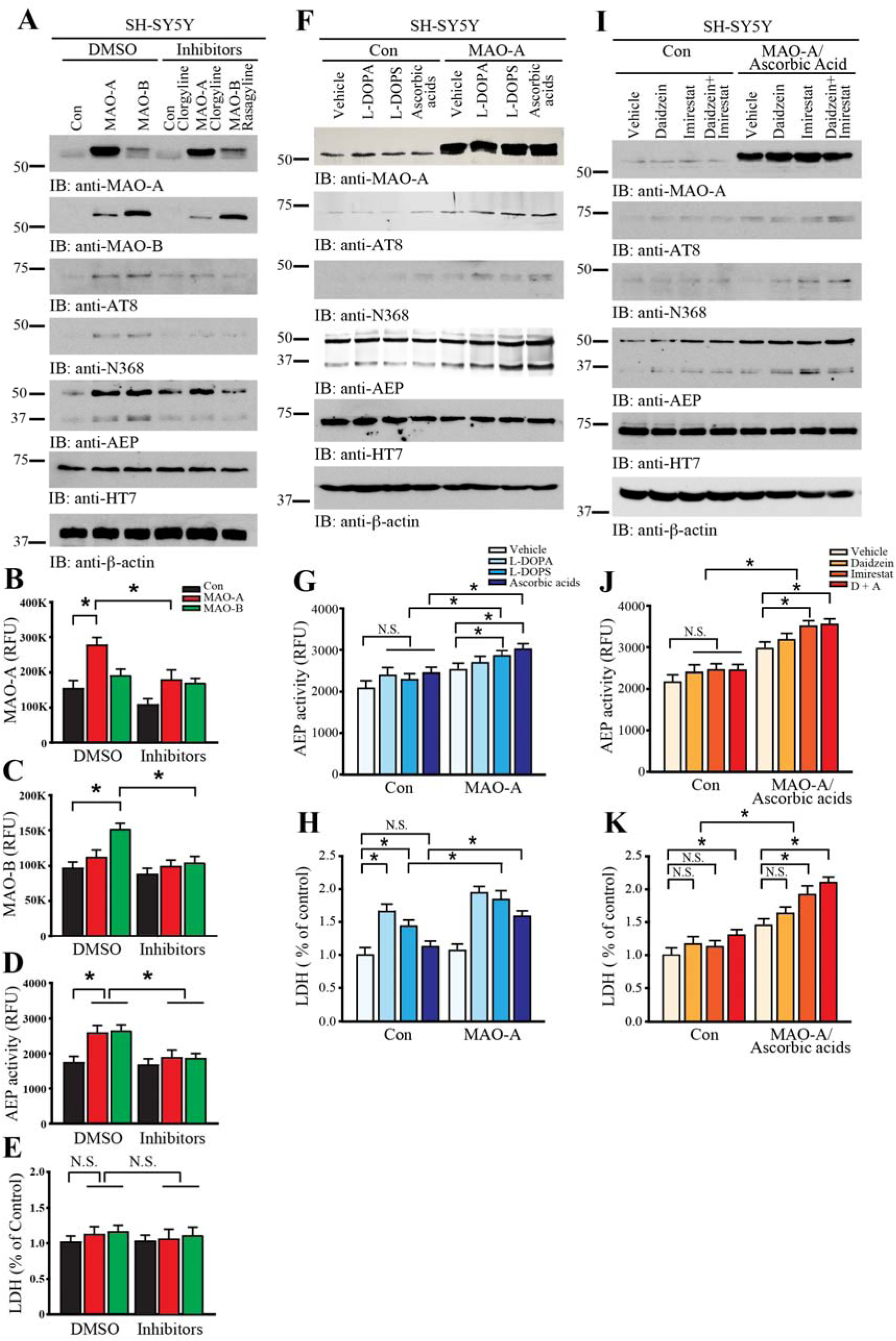
DOPEGAL induces Tau neurotoxicity in SH-SY5Y cells. **A.** SH-SY5Y cells were transfected with MAO-A or MAO-B, followed by treatment with the MAO-A inhibitor clorgyline (10 μM) or the MAO-B inhibitor rasagyline (10 μM). Western blot analysis showed that MAO overexpression induced AEP activation, Tau phosphorylation, and Tau cleavage, and MAO inhibitors blocked these events. **B-D**. The activation of MAO-A (**B**), MAO-B (**C**), and AEP (**D**) was confirmed by enzymatic assay. **E**. LDH assay showed that MAO-A or MAO-B overexpression in SH-SY5Y cells did not induce cell death on their own. Data are shown as mean ± SEM. N=3 per group. * p<0.05. **F**. SH-SY5Y cells were transfected with MAO-A, followed by treatment with L-DOPA (200 μM), L-DOPS (200 μM), or Ascorbic acid (500 μM). Immunoblotting showed that MAO-A overexpression induced AEP activation, Tau phosphorylation, and Tau cleavage in the presence of NE precursors. **G & H**. AEP activity and cell death were increased by L-DOPA, L-DOPS, or Ascorbic acid with MAO-A overexpression. Data are shown as mean ± SEM. N=3 per group. * p<0.05. **I.** SH-SY5Y cells were transfected with MAO-A and treated with Ascorbic acid (500 μM). And then cells were treated with vehicle, the aldehyde dehydrogenase inhibitor Daidzein (10 μM), the aldehyde reductase inhibitor Imirestat (1 μM), or their combination (D+I) for 24 h. Western blot analysis showed that inhibition of DOPEGAL metabolism induced AEP activation, Tau phosphorylation, and Tau cleavage following facilitation of NE production and MAO-A oxidation. **J & K**. AEP activity and cell death were increased by the treatment of Daidzein, Imirestat, or inhibitors combination with MAO-A overexpression and ascorbic acid exposure. Data are shown as mean ± SEM. N=3 per group. * p<0.05.

**Supplemental Figure 5.**
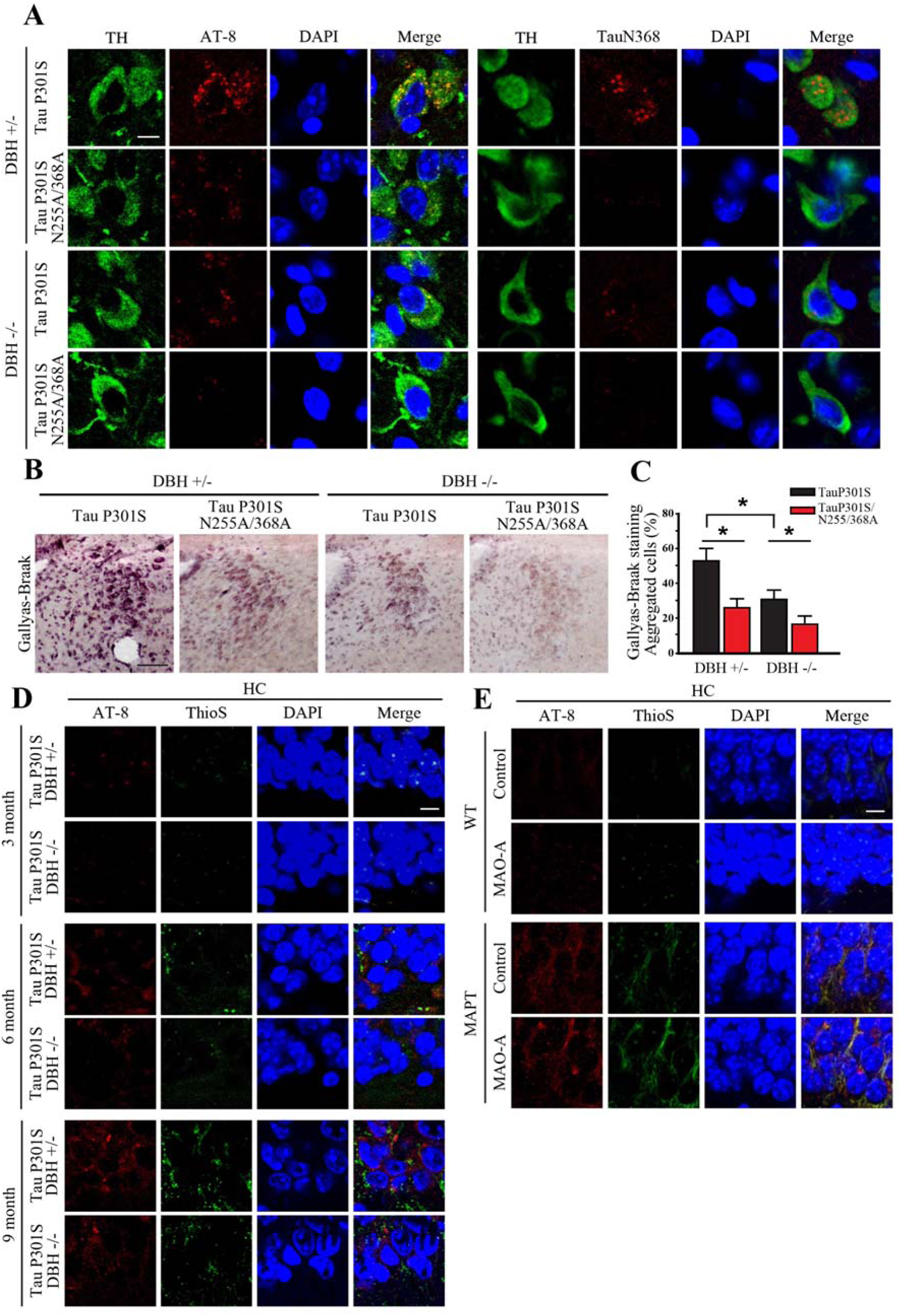
Tau cleavage by AEP is necessary for Tau pathology progression in LC region. **A.** Tau P301S or AEP cleavage-resistant Tau P301S/N255/368A virus was injected into the LC of DBH +/− or DBH −/− mice. Immunofluorescent co-staining showed that Tau N368 and AT8 were increased in the LC of Tau P301S-injected mice 3 months after viral injection, and that these effects were inhibited by DBH knockout or blockade of Tau cleavage. Scale bar = 20 μm. **B.** Gallyas-Braak staining showed that Tau aggregation following infection of Tau P301S was inhibited in DBH −/− mice or by preventing Tau P301S cleavage. Scale bar = 100 μm. **C**. Quantification of aggregated cells from Gallyas-Braak stained images. Data are shown as mean ± SEM. N=4 per group. * p<0.05. **D.** Representative images of immunofluorescent staining for AT8 (red), ThioS (green), and DAPI (blue) in the HC 3, 6, or 9 months following viral injection. **E.** Control or MAO-A AAV were injected into the LC of wild-type (WT) or MAPT transgenic mice, and Tau pathology was assessed 2 months later. Shown are representative images of immunofluorescent staining for AT8 (red), ThioS (green), and DAPI (blue). Scale bar = 20 μm.

**Supplemental Figure 6.**
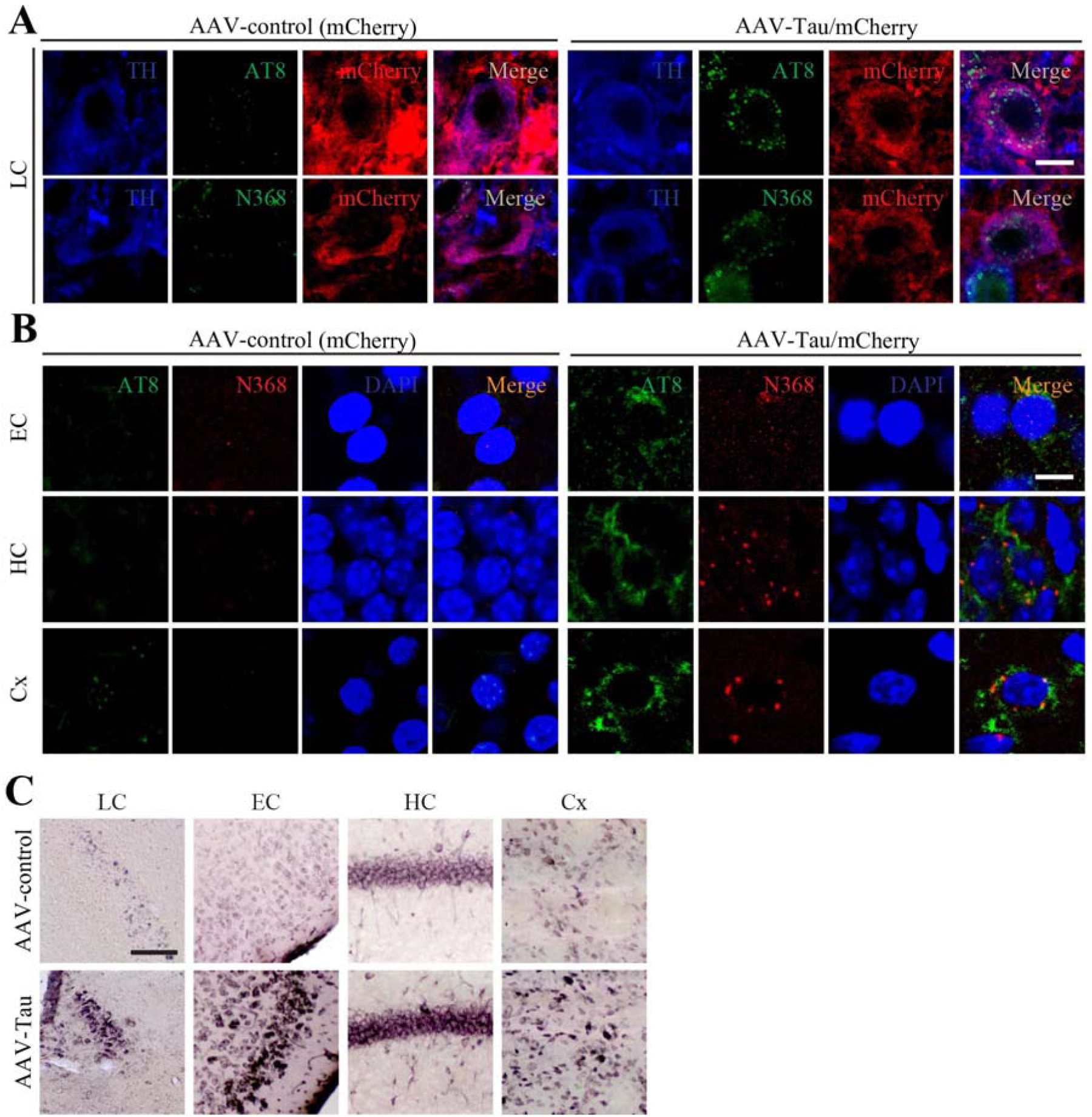
Viral-mediated Tau expression in the LC drives propagation of pathology to the forebrain in 3xTg transgenic mice. LC-specific AAV-PRSx8-Tau + AAV-PRSx8-mCherry or AAV-PRSx8-mCherry alone were injected into the LC of 3xTg mice, and cognition and pathology were assessed 3 months later. **A**. Representative images of immunofluorescent staining of TH with AT8/ mCherry or N368/mCherry shows that AAV-Tau infection induces Tau phosphorylation and cleavage in LC. Scale bar = 20 μm. **B**. Representative images of immunofluorescent staining of TH with AT8/ mCherry or N368/mCherry in Entorhinal cortex (EC), Hippocampus (HC), and Cortex (Cx) sections. Scale bar = 20 μm. **C**. Gallyas-Braak staining shows that Tau aggregation induced by Tau is detected in LC, EC, HC, and Cx. Scale bar = 100 μm.

**Supplemental Figure 7.**
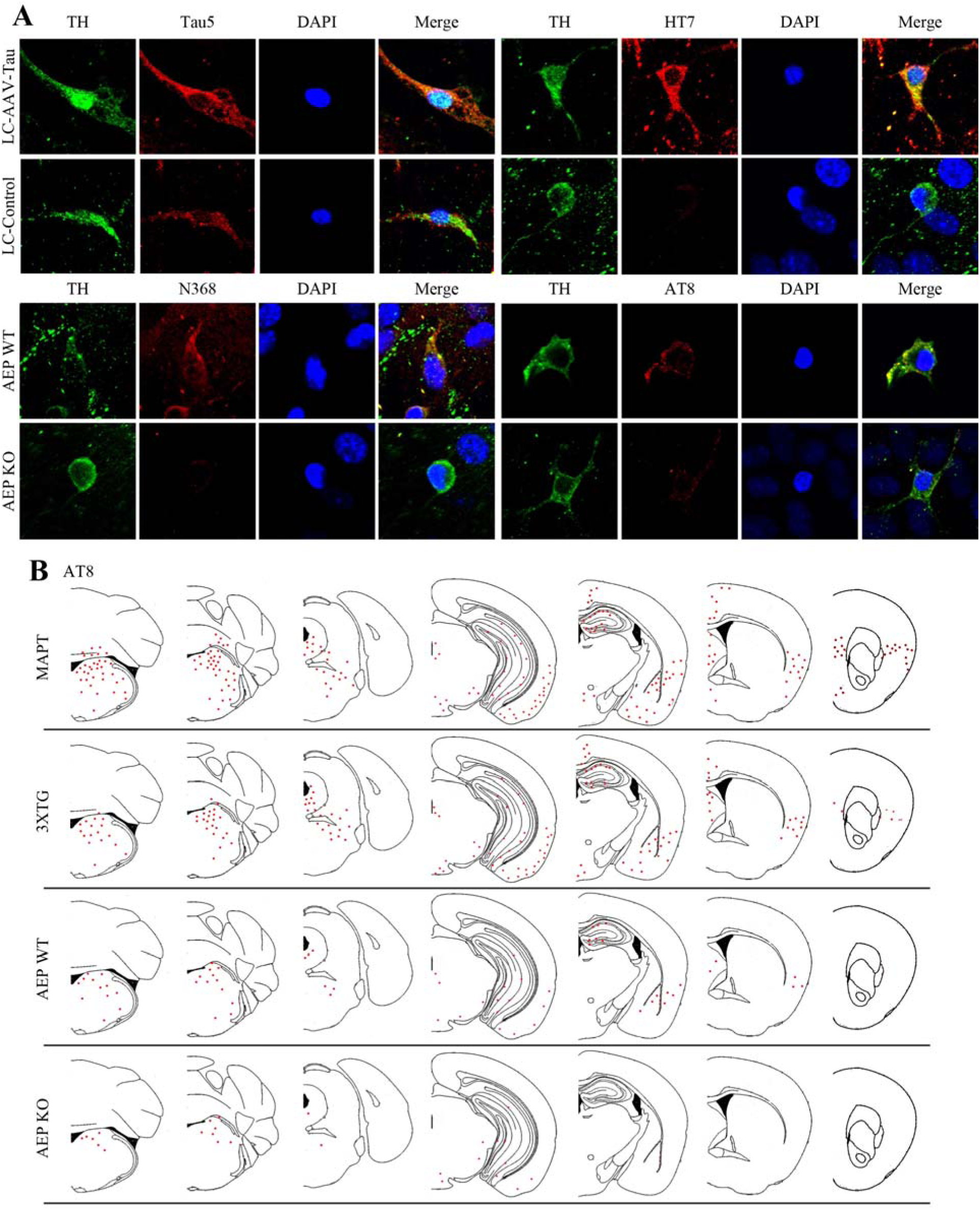
Development of Tau pathology in the LC and propagation to the forebrain is dependent on AEP. **A.** Primary LC neurons were prepared from neonatal AEP +/+ or AEP −/− mice and infected with LC specific AAV-PRSx8-Tau or control virus. Upper panels show representative immunofluorescent images for TH (green), total Tau or human Tau (red) in AEP +/+ neurons infected with AAV-PRSx8-Tau or control virus. Lower panels show representative immunofluorescent images for TH (green), AT8 or Tau N368 (red), and DAPI (blue) in AEP +/+ or AEP −/− neurons infected with AAV-PRSx8-Tau. LC neurons from AEP −/− mice were resistant to Tau phosphorylation and cleavage. Scale bar = 20 μm. **B.** Schematic summary maps of AT8 immunohistochemistry showing Tau pathology spreading from the LC to different brain regions in MAPT, 3XTG, AEP +/+, and AEP−/− mice.

